# Nuclear Export of *Drosophila* PERIOD contributes to temperature compensation of the circadian clock

**DOI:** 10.1101/2021.10.25.465663

**Authors:** Astrid Giesecke, Peter S Johnstone, Angelique Lamaze, Johannes Landskron, Ezgi Atay, Ko-Fan Chen, Deniz Top, Ralf Stanewsky

## Abstract

Circadian clocks are self-sustained molecular oscillators controlling daily changes of behavioral activity and physiology. For functional reliability and precision the frequency of these molecular oscillations must be stable at different environmental temperatures, known as ‘temperature compensation’. Despite being an intrinsic property of all circadian clocks, this phenomenon is not well understood at the molecular level. Here we use behavioral and molecular approaches to characterize a novel mutation in the *period* (*per*) clock gene of *Drosophila melanogaster*, which alters a predicted nuclear export sequence (NES) of the PER protein. We show that this new *per^I530A^* allele leads to progressively longer behavioral periods and clock oscillations with increasing temperature in both clock neurons and peripheral clock cells. While the mutant PER^I530A^ protein shows normal circadian fluctuations and post-translational modifications at cool temperatures, increasing temperatures lead to both, severe amplitude dampening and hypophosphorylation of PER^I530A^. We further show that PER^I530A^ displays reduced repressor activity at warmer temperatures, presumably because it cannot inactivate the transcription factor CLOCK (CLK). With increasing temperatures nuclear accumulation of PER^I530A^ within clock neurons is increased, suggesting that PER is normally exported out of the nucleus at warm temperatures. Consequently, downregulating the nuclear export factor CRM1 also leads to temperature-dependent changes of behavioral rhythms. In summary, our results suggest that the PER NES and the nuclear export of clock proteins play an important role in temperature compensation of the *Drosophila* circadian clock.

## Introduction

Circadian clocks allow organisms to anticipate the daily changes of their environment, such as daily fluctuations of light intensity and temperature. They enable animals to restrict their behavioral and physiological activities to species-specific advantageous times of day and night, increasing their overall fitness. Circadian clocks are composed of negative molecular feedback loops, within which clock gene products oscillate in abundance and subcellular localization with a 24-h period (1). In *Drosophila melanogaster*, expression of the clock genes *per* and *timeless* (*tim*) is mediated by the transcription factors CLK and CYCLE (CYC). PER and TIM proteins accumulate in the cytoplasm before they translocate to the nucleus, bind to the CLK/CYC dimer and repress their own transcription. Eventually, PER and TIM proteins are degraded, allowing CLK and CYC to start a new round of the cycle (1). Posttranslational modifications regulate this process to maintain the period length of the molecular oscillations at ∼ 24 h. For example, cytoplasmic PER and TIM phosphorylation by the kinases DBT, CK2 and SGG determines timed nuclear translocation (2). In addition, hyperphosphorylated nuclear PER coincides with hyperphosphorylated, inactive CLK, and the DOUBLETIME (DBT) kinase mediates phosphorylation of both proteins within the same complex leading to transcriptional repression (3, 4). In the so called PER ‘phosphotimer’ the NLK kinase NEMO initially phosphorylates PER S596, which stimulates phosphorylation of S589, S585 and T583 by the CK1ε kinase DOUBLETIME (DBT). All these residues belong to the *per^Short^* phosphocluster, and their phospho-occupancy somehow delays DBT-dependent PER phosphorylation at other residues, most importantly PER S47 (5). Since phosphorylation at PER S47 enables binding of the F-box protein SLIMB and subsequent degradation of PER (6), these consecutive phosphorylation events regulate temporal PER stability and thereby period length of the circadian clock (5). Based on the involvement kinases and phosphatases and other post-translational modifications in clock regulation it could be expected that increasing temperature leads to faster enzyme kinetics and therefore shorter periods of molecular oscillations and vice versa. However, circadian clocks are temperature compensated, meaning periods of circadian clock oscillations do not change across a wide range of temperatures (e.g., between ∼15°C and 29°C in fruit flies) (7). Although temperature compensation is a hallmark of all circadian clocks (8, 9), its molecular etiology is still not well understood. Circadian clocks can be synchronized by temperature cycles, which suggests that they are temperature-responsive with compensatory mechanisms (10). This compensation is traditionally explained with the assumption that temperature-mediated changes of different rates across the circadian clock cancel each other out (11). This model is less likely, given that any mutation in circadian genes would yield a temperature-sensitive circadian clock.

Temperature compensation could also be achieved by considering the circadian clock as two coupled oscillators with complementary temperature dependence of period length (10), or as one oscillator composed of biochemical reactions with opposite temperature coefficients (12). This would require different clocks within one organism to have distinct regulatory mechanisms (13). The circadian clock of *Drosophila* consists of multiple coupled oscillators that vary in their intrinsic period length (e.g. (14)), but it is not known how they behave at different temperatures. Subsets of these oscillators, such as the so-called small lateral neurons (s-LNv), sustain free-running behavioral rhythms in constants darkness (DD) (15), while others such as the Dorsal Neurons 1 - 3 (DN1, DN2, and DN3) are unable, despite sustaining molecular rhythms in DD (16, 17). It is therefore conceivable that the different clock neuronal subsets also differ in their ability to maintain temperature-compensated molecular oscillations.

Another possible mechanism for temperature compensation, is that each step in the clock is individually temperature compensated (18). For example, timed nuclear accumulation and timed nuclear degradation of PER/TIM would each be temperature compensated. Some alleles of *per* and *tim* that are defective in nuclear accululation of the PER/TIM complex also affect temperature compensation. *per^L^* flies (19) exhibit an increase of behavioral period by four hours from 18°C (27 h) to 29°C (31 h), which correlates well with the ∼4-hour delay of PER^L^ nuclear entry (20, 21). This delay can be explained by the weaker interaction of PER^L^ with TIM with increasing temperatures, because TIM stabilizes PER and PER^L^ therefore accumulates even slower at higher temperatures (22).

Similarly, the *tim^blind^* allele increases behavioral period by four hours in a temperature-dependent manner from 18°C (24 h) to 29°C (28 h) (23). This, and several other new *tim* alleles map to nuclear export signals (NESs) throughout TIM and also show similar temperature compensation defects (23). Indeed, TIM and PER are subject to active Cargo Protein and Chromosome Maintenance 1 (CRM1)-mediated nuclear export in Drosophila larvae (24). With TIM export, this mechanism relies on CKII-dependent phosphorylation of residue S1404 near a putative NES, which reduces the interaction between TIM and CRM1, and retains normal nuclear levels of TIM and regulation of CLK activity (3). In mammals, mPER1, mPER2, and mPER3 proteins also contain canonical leucine-rich NES motifs, which in cell culture shuttle PER proteins from the nucleus to the cytoplasm in order to target them for degradation (25, 26), but their roles in temperature compensation have not been examined.

While it seems clear from these studies that nuclear export of clock proteins plays a role in circadian clock function in both mammals and fruit flies, its role in temperature compensation has not been examined. Here, we generated a point mutation (*per^I530A^*) in the single leucine-rich putative PER NES, which is highly conserved to the mPER NES (Figure 1A) (25,27,28) to systematically study the role of the PER NES in the context of temperature compensation. *per^I530A^* mutant flies show substantial temperature-dependent increase of their behavioral period, which is correlates well with longer molecular oscillations in several subsets of clock neurons and in peripheral clock cells. PER^I530A^ shows temperature-dependent alterations of posttranscriptional modifications, accompanied by reduced ability to function as a transcriptional repressor of CLK. Finally, enhanced nuclear PER^I530A^ accumulation with increasing temperatures, as well as temperature-dependent period lengthening of flies with reduced CRM1 expression strongly implicate nuclear export of PER as an important mechanism for temperature compensation in Drosophila. The knockdown of CRM1 weakens the observed temperature-compensation defect of *per^I530A^*, suggesting that nuclear export of other clock proteins may also be affected, and implicating nuclear export as a key mechanism in establishing temperature compensation.

**Figure 1.**
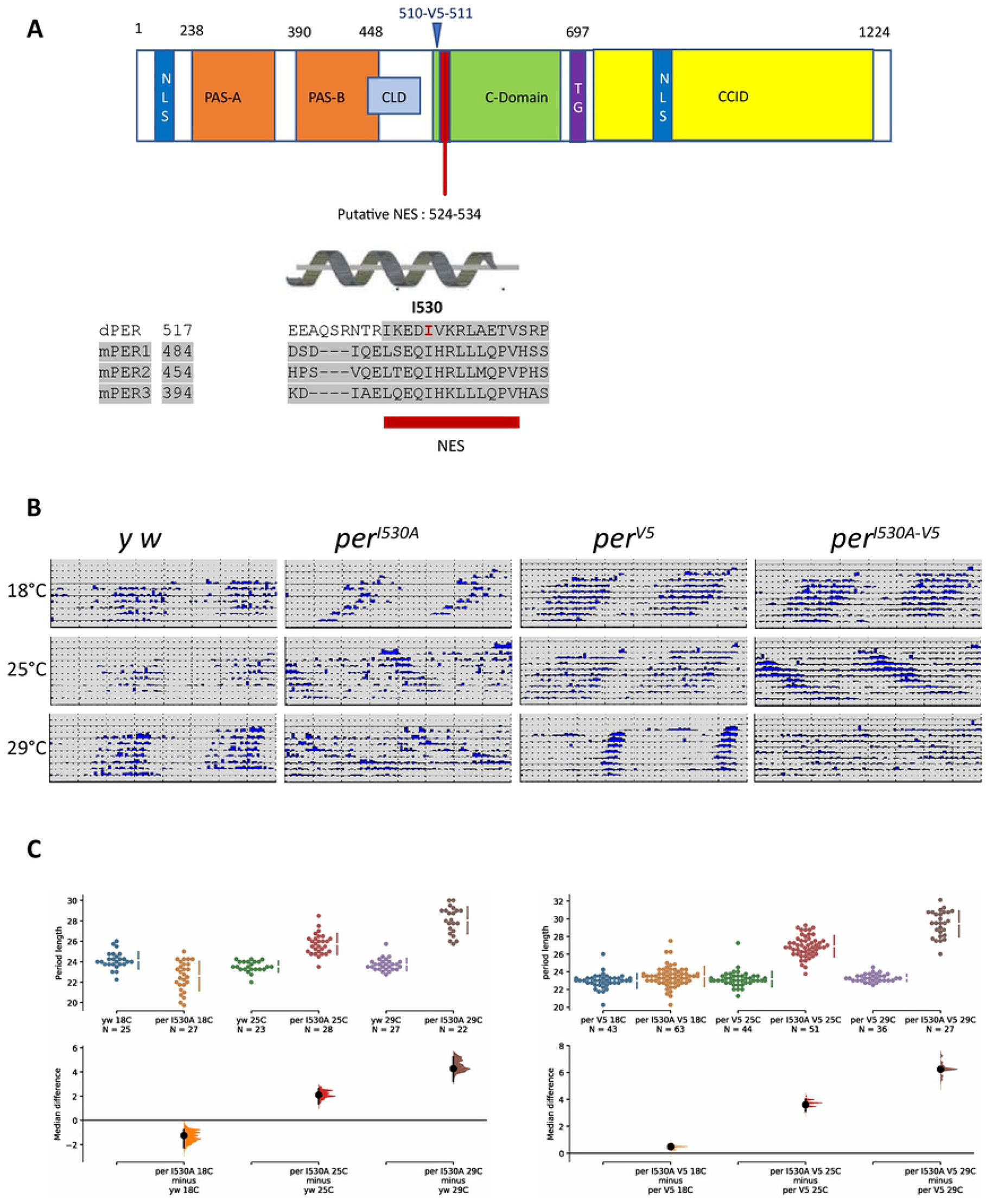
Effect of *per^I530A^* on temperature compensation: (A) Cartoon of the PER structure (amino acids 1–1,224) showing the different functional domains including the putative NES and the position of the inserted V5 tag. A magnification of the NES region between residues 510-525 is shown below including a sequence alignment of the putative NES domain from DmPER and mPER1-3. Grey colored regions mark the conserved hydrophobic sites. The position of the Isoleucine 530 (mutated to Alanine in *per^I530A^*) is shown in red. The alpha-helix formed by the NES is depicted above the sequence alignment. (B) Effect of *per^I530A^* on free-running locomotor activity analyzed at three different temperatures. Flies were kept for 3-4 days in LD followed by an additional week in DD at the constant temperatures indicated on the left side of each panel. For controls (*y w* and *per^V5^*) and for *per^I530A^* flies (with or without V5 tag) individual actograms of the LD and DD part of the experiments are shown. (C) Period estimates and statistical analysis of individual flies. To analyze the distribution of period values at the different temperatures estimation statistics (ES) was applied. The top part shows the individual period data points obtained with the standard error plotted as a line to the right, with the gap indicating the average. The lower part summarizes the median difference between *per^I530A^* and control. The colored curves represent the median difference distribution of the values, and the black vertical line indicates the 95% confidence interval (see Materials and Methods for details).

## Results

### The *per^I530A^* mutant causes circadian period lengthening with increasing temperatures

To analyze the function of the nuclear export signal (NES) in the *Drosophila* Period (PER) protein, we introduced a single amino acid change into the NES by replacing the conserved isoleucine at position 530 with alanine (PER^I530A^). To do this, the endogenous *per* gene was replaced with the mutated *per^I530A^* allele by CRISPR/Cas9 mediated homologous recombination (Figure 1A and Methods). To aid detection of wild type and mutated PER^I530A^ we also generated flies where both proteins are endogenously tagged with the V5 epitope (*per^V5^* and *per^I530A-V5^* respectively, Figure 1A and Methods). To investigate if the *per^I530A^* allele influences circadian clock function and behavioral rhythms, we analyzed locomotor activity in constant darkness at 25°C (DD25). While *y w* control flies showed the typical robust activity rhythms with period values (τ) of 23.5 hr, τ values of the *per^I530A^* flies were > 2hr longer (25.9 hr) (Figure 1B, C, Table 1). Similarly, *per^V5^* flies have τ values of 23.2 hr compared to 26.7 hr in *per^I530A-V5^* flies (Figure 1B, C, Table 1). To test if the PER NES is also important for temperature compensation, we tested *per^I530A^* flies at different temperatures (DD18 and DD29). Indeed, *per^I530A^* flies shortened their period by ∼3 hr to 23.0 hr at 18°C and lengthened it to 27.9 hr at 29°C (Q10=0.84), while *y w* controls showed almost no change (Q10=0.99) (Figure 1B, C, Table 1). Similar results were obtained with the V5-tagged flies: while *per^V5^* flies exhibit stable τ values around 23 hr from 18°C to 29°C (Q10=1.0), the *per^I530A-V5^* flies increase τ from 23.5 hr at 18°C to 29.4 hr at 29°C (Q10=0.82) (Figure 1B, C, Table 1). The results unequivocally show that specific mutation of the PER NES leads to a temperature-dependent lengthening of τ, pointing to temperature over-compensation of the circadian clock.

**Table 1:** Free-running behavioral rhythms of *per^I530^* mutants and controls

| Genotype | Temp (°C) | n | Tau (hr ± SEM) | RS ± SEM | % rhythmic |
| --- | --- | --- | --- | --- | --- |
| *y w* | 18 | 30 | 24.1 ± 0.2 | 3.0 ± 0.2 | 83.3 |
|  | 25 | 32 | 23.5 ± 0.1 | 2.6 ± 0.2 | 71.9 |
|  | 29 | 29 | 23.7 ± 0.1 | 2.8 ± 0.2 | 93.1 |
| *per*<sup>L530A</sup> | 18 | 38 | 23.0 ± 0.4 | 2.2 ± 0.2 | 71.1 |
|  | 25 | 35 | 25.9 ± 0.3 | 2.4 ± 0.3 | 80.0 |
|  | 29 | 37 | 27.9 ± 0.3 | 1.8 ± 0.1 | 59.5 |
| *per*<sup>V5</sup> | 18 | 44 | 23.0 ± 0.1 | 3.4 ± 0.2 | 97.7 |
|  | 25 | 47 | 23.2 ± 0.1 | 3.5 ± 0.2 | 93.6 |
|  | 29 | 43 | 23.3 ± 0.1 | 3.4 ± 0.3 | 83.7 |
| *per*<sup>L530A-V5</sup> | 18 | 78 | 23.5 ± 0.1 | 3.0 ± 0.2 | 80.8 |
|  | 25 | 68 | 26.7 ± 0.2 | 2.3 ± 0.2 | 75.0 |
|  | 29 | 47 | 29.4 ± 0.3 | 1.9 ± 0.2 | 57.4 |

### The *per^I530A^* mutation leads to temperature dependent molecular period lengthening within central clock neurons and peripheral clock cells

Behavioral rhythms in constant darkness are driven by molecular oscillations of clock gene expression within circadian pacemaker neurons in the fly brain. It is therefore expected that temperature-dependent period-lengthening also occurs in clock neurons of *per^I530A^* mutant flies. To test this, and to see if temperature compensation is affected equally in all clock neurons, we applied a novel, cell-type restricted real-time luciferase assay (<u>L</u>ocally <u>A</u>ctivatable <u>B</u>ioluminescence or LABL). Central to LABL is a *period-luciferase* transgene, in which *luciferase* expression is compromised by a mCherry reporter gene followed by stop codons. This cassette is inserted between the *period* promoter and the *luciferase* coding sequences, and flanked by FRT recombination sites (Figure S1A). *period-*driven *luciferase* expression can therefore be induced and restricted to subsets of *per* expressing cells using Gal4 driven expression of UAS-FLP (Figure S1A). Details of this technique are described in an accompanying paper (Johnstone et al 2021, this issue). Using LABL we first compared *period-luciferase* expression in different sets of clock neurons using the *Pdf-gal4* (PDF positive s-LNv and l-LNv), *DvPdf-gal4* (PDF positive s-LNv and l-LNv, 5^th^ s-LNv, 4 LNd) and *R18H11-gal4* (subset of the DN1p) at three different temperatures in wild type flies. Bioluminescence of 15 male flies per genotype housed in a customized arena was measured using the LumiCycle 32 Luminometer (Actymterics) for 9 days in constant darkness at either 18°C, 25°C, or 29°C (see Materiala and Methods and Johnstone et al 2021, this issue, for details). In wild type flies, all driver lines generated robust bioluminescence oscillations with stable period lengths close to 24 hr at all temperatures (Figure 2A, C, S1B), indicating that all subsets of clock neurons are temperature compensated (cf. (29)). In contrast, the same analysis in the background of the *per^I530A^* mutation revealed a significant linear period-lengthening from 18°C to 29°C for *Pdf* and *R18H11-gal4* driver lines and between 18°C and 25°C for *DvPdf-gal4* (Figure 2A, C, S1B) (for unknown reasons the genotype involving *per^I530A^* and *DvPdf-gal4* died at 29°C). At the lower temperature, bioluminescence oscillations showed a period close to 24 hr, while at 25°C it was lengthened from 26-28 hr, and further to 30-32 hr at 29°C, depending on the Gal4 driver used to activate LABL (Figure 2D, S1B). Therefore, the period length of the molecular oscillations and behavioral rhythms are equally affected by the *per^I530A^* mutation, showing normal values at 18°C and temperature-dependent period lengthening. As a control, we also analyzed *period-luciferase* expression in *per^L^* flies, which have a period of 27.1 hr at 18°C, increasing to 29.6 hr at 25°C and 31.4 hr at 29°C (20). Again, using *Pdf-gal4* to drive LABL, the molecular oscillations closely matched the behavioral ones, with periods of 27-28 hr at 18°C lengthening to 29-30 hr at 25 °C, and 32 hr at 29°C (Figure S1C). We conclude that LABL is suitable for reliable period estimation within clock neurons at different temperatures (see also accompanying paper by Johnstone et al).

**Figure 2.**
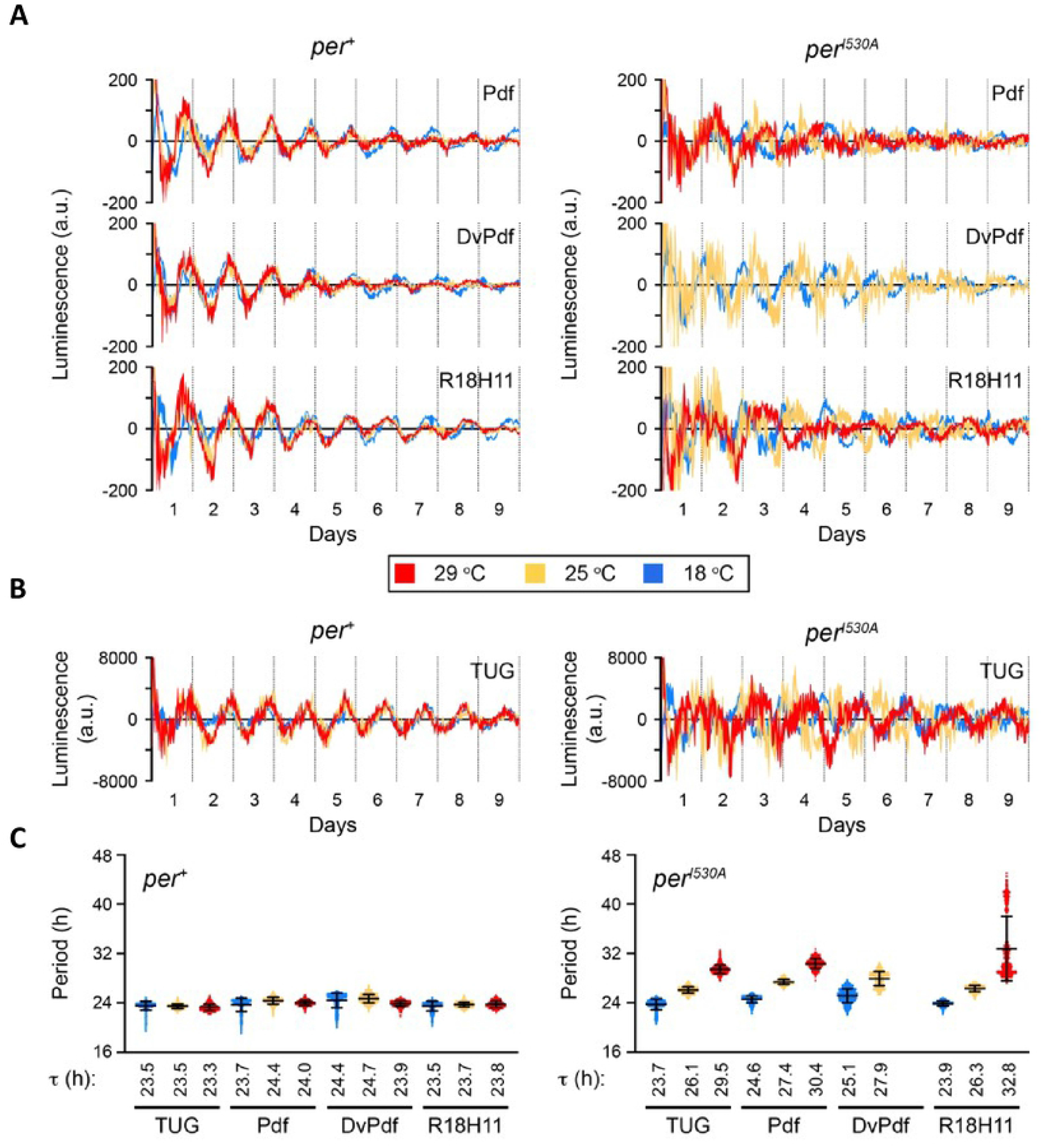
*per^I530A^* affects temperature compensation in different neuronal clocks: (**A**) of Bioluminescence oscillations measured from LABL flies activated by the indicated circadian neuronal drivers (*Pdf-Gal4, DvPdf-Gal4* and *R18H11-Gal4*), in a wild type (*per*^+^) or mutant (*per^I530A^*) genetic background, plotted over time. Colors represent mutant flies monitored at different temperatures: 18 °C (blue), 25 °C (yellow) or 29 °C (red). Line thickness represents the mean of two experiments, +/- SEM. (**B**) Bioluminescence oscillations measured from LABL flies activated by the universal circadian driver *tim-UAS-Gal4*. Figure details are as described in panel A. (**C**) Average periods of *per-luc* oscillations in different neurons. Indicated Gal4 drivers were used to activate LABL in wild type (*per*^+^) or mutant (*per^I530A^*) flies. Colors represent different temperatures as described in panel A. Bar represents the mean, +/- SD. Mean values of period (τ) are denoted underneath the graph.

To determine whether temperature-dependent defects of the *per^I530A^* allele also extend to peripheral clocks, we next used LABL to measure *period-luciferase* expression in all clock cells. For this, we applied the *tim-UAS-Gal4* driver expressed in all clock cells (30) Johnstone et al 2021, this issue). Although the *tim-UAS-Gal4* driver will also activate *period-luciferase* expression in clock neurons, the vast majority of bioluminescence expression emanates from peripheral clock cells in abdominal tissues and in the head (Johnstone et al 2021, this issue), as previously shown for *period-luciferase* transgenes expressed in all clock cells (31). As expected, the bioluminescence levels emanating from *tim-UAS-Gal4* LABL flies were drastically (40-50 fold) increased compared to the three clock neuronal drivers, reflecting expression in peripheral clock cells (Figure 2B). Similar to the results obtained for the clock neurons, oscillations in peripheral clock cells in wild type flies were temperature-compensated, with stable periods close to 24 hr at all three temperatures (Figure 2B, C, S1B). Moreover, when analyzed in a *per^I530A^* and *per^L^* mutant background, *tim-UAS-Gal4* driven *period-luciferase* oscillations showed the same temperature-dependent period-lengthening as observed with the neuronal drivers (Figure 2B, C, S1B, C). Therefore, the *per^I530A^* mutation has a similar effect on both central and peripheral clocks, indicating a common mechanism of temperature compensation.

### The *per^I530A^* mutation affects PER and TIM stability and post-translational modification at higher temperatures during LD

The temperature-dependent period-lengthening of *per^I530A^* flies could be due to decreased PER^I530A^ stability at higher temperatures. To test this, we performed Western blots of total head protein extracts of control (*per^V5^*) and *per^I530A-V5^* flies raised at different temperatures and collected every six hours during one day in LD (ZT2, ZT8, ZT14, ZT20). To see how wild type PER levels behave at different temperatures throughout the day we first analyzed *per^V5^* flies at 18°C, 25°C and 29°C (Figure 3). As expected, we observed robust PER oscillations at all three temperatures, with trough levels at ZT8 and peak expression from ZT20 to ZT2 (Figure 3A) (32). At 18°C PER levels appear somewhat reduced from ZT2 to ZT14 compared to the higher temperatures, but the differences were not significant (Figure 3A). We also did not detect obvious differences in the migration speed of PER, which is indicative of its phosphorylation status (33), except that at ZT2 during 29°C PER migrates faster compared to the cooler temperatures and therefore appears hypophosphorylated. These results show that wild type PER levels as well as rhythmic changes in abundance and phosphorylation are well compensated against changes of ambient temperature.

**Figure 3.**
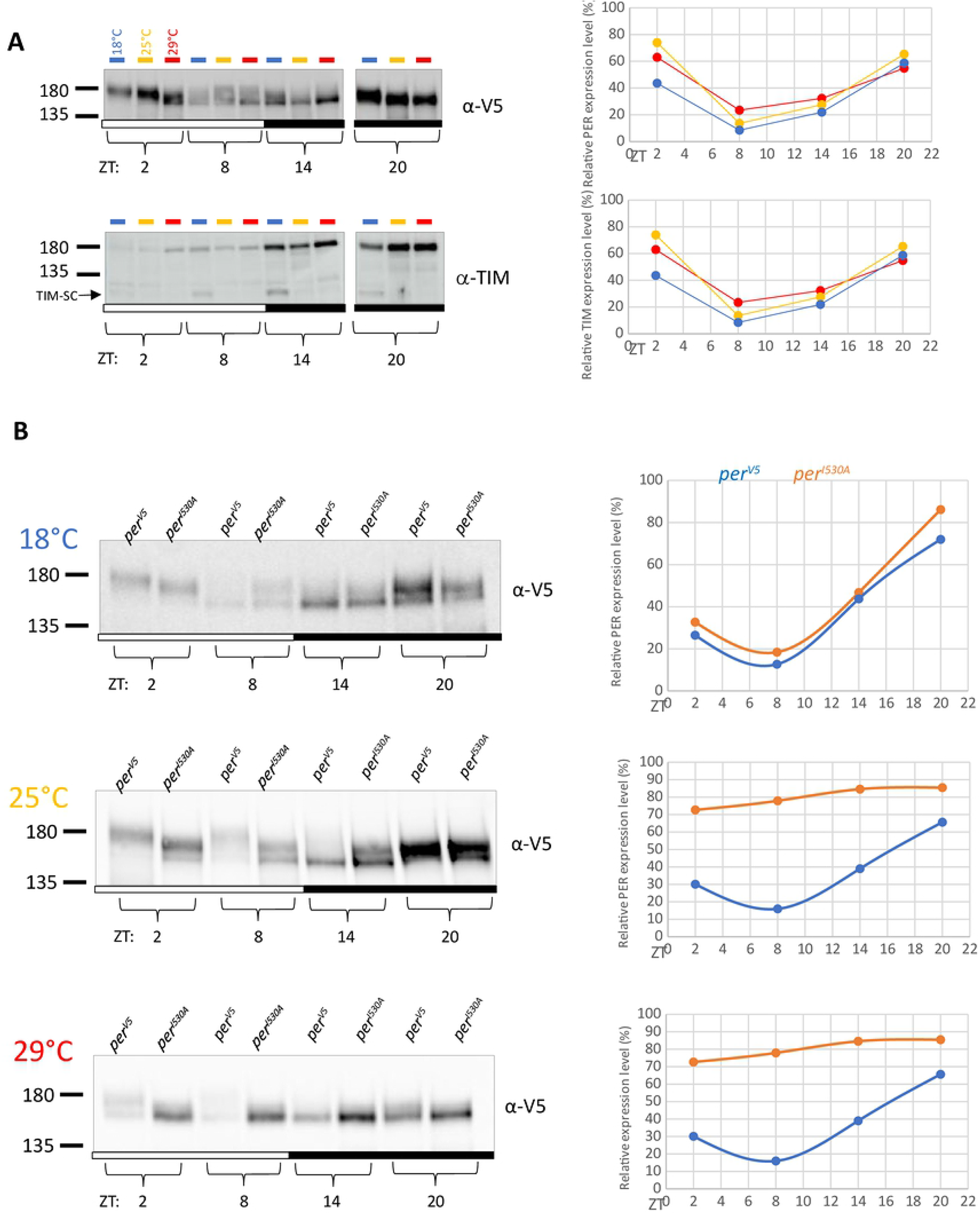
*per^I530A^* causes temperature-dependent defects in oscillation and posttranslational modification of PER. **(A)** Expression of wild type PER carrying the V5 tag was analyzed at different temperatures. Flies were kept for 3 days in LD at three constant temperatures (18°C, 25°C and 29°C). Protein extracts from adult fly heads were prepared at the indicated Zeitgeber Times (ZT) as described in Materials and Methods. Left panel: western blots comparing PER and TIM temporal expression profiles at different temperatures. Blots were incubated with anti-V5 or anti-TIM antibodies, respectively. Right panel: Relative expression levels were plotted as line graphs showing the PER or TIM protein oscillations during a 24 h day at the three different temperatures. Error bars indicates SEM. **(B)** PER expression in *per^V5^* and *per^I530A-V5^* was compared at the indicated time points and temperatures. Left panels: Protein extract from wild type mutant flies were loaded next to each other for each time point to allow direct comparison. Right panels: Relative expression levels were plotted as line graphs for *per^V5^* and *per^I530-V5^* flies. Error bars indicate SEM.

Next we compared PER and PER^I530A^ expression at different temperatures. At 18°C no obvious differences in protein amounts or mobility could be observed during the four time points we analyzed (Figure 3B, top panel). In contrast, at 25°C an amplitude reduction of PER^I530A^ cycling due to increased trough levels at ZT8 was apparent (Figure 3B, middle panel). In addition, there was a striking effect on mobility at ZT2 and ZT8. At these time point wild type PER appears mainly in its slow migrating form, which is not observable for PER^I530A^. Instead the faster migrating form is still present at ZT8, indicating higher stability of PER^I530A^ due to incomplete post-translational modifications and degradation. This could ultimately result in the lower-amplitude protein cycling we observed (Figure 3B, middle panel). Strikingly, at 29°C PER^I530A^ oscillations were completely abolished and presumably hypophosphorylated protein was present at peak levels at all time points analyzed (Figure 3B, lower panel). To summarize, in parallel to the behavioral and luciferase observations, PER^I530A^ protein behaves normal at 18°C, but both, protein cycling and the phosphorylation pattern become increasingly aberrant with rising temperature. Our results suggest that a temperature-dependent phosphorylation defect underlies the increased stability and reduced oscillation of the PER^I530A^ protein at higher temperatures.

Next, we examined if the *per^I530A^* mutation had similar effects on the clock protein TIM. First, we again checked if TIM expression varied in wild type in a temperature-dependent manner. We did not detect any differences in levels or migration properties, except for the expected detection of the shorter, cold-induced TIM-SC protein at 18°C (Figure 3A, lower panel) (34, 35). As expected from the results with PER, wild type TIM levels and phosphorylation patterns are therefore well compensated against temperature changes. Comparison of TIM between wild type and *per^I530A^* mutants revealed only mild differences at the warmer temperatures, while at 18°C they were essentially identical at all time points (Figure S2). At 25°C TIM levels in *per^I530A^* were slightly increased at ZT2 and ZT8, and this effect was further pronounced at 29°C (Extended Data Figure 3). Nevertheless, even though the amplitude is reduced, TIM levels still oscillated. We were not able to detect any differences in TIM migration between wild type and *per^I530A^*, but phosphorylation-dependent migration differences are generally not as pronounced for TIM compared to PER (compare Figure 2A, B with S2).

### The *per^I530A^* mutation affects PER and TIM stability and post-translational modification at higher temperatures during DD

Because the temperature-dependent effects on behavioral period length are necessarily determined in constant darkness, we wanted to analyze the effects of the *per^I530A^* allele on PER and TIM expression also in DD. PER and TIM isolated from total fly heads mainly reflect expression in the photoreceptor cells, which contain peripheral clocks that dampen out rapidly in DD (16). We therefore analyzed PER and TIM levels during the first day in DD after LD entrainment from both wild type and *per^I530A^* flies at different temperatures. At 18°C, both PER and TIM cycled with similar amplitude in mutants and controls, although both proteins appeared more stable in *per^I530A^* (Figure S3). At 25°C and 29°C weak PER and TIM oscillations were observed in wild-type, whereas in the *per^I530A^* mutants both proteins were at high constitutive levels (Figure S3). Similar as in LD, PER^I530A^ protein showed normal daily migration changes at 18°C, while at 25°C and 29°C the slow migrating, hyperphosphorylated forms that normally occur in the subjective morning (CT2 and CT8) were absent and instead high levels of faster migrating hypophosphorylated PER^I530A^ were visible (Figure S3).

In order to track the effects of *per^I530A^* on PER oscillations with high temporal resolution, we turned to the *BG-luc* reporter, a *period-luciferase* transgene encoding for ∼2/3rds of the PER protein. This *per* transgene does not rescue behavioral and molecular arrhythmicity induced by the *per^01^* allele, but it accurately reports PER protein cycling in a wild-type background (31), and is therefore suitable to determine the effects of *per* alleles on PER protein expression. Bioluminescence rhythms emanating from individual adult male *BG-luc* flies were measured for four days in LD followed by three days in DD at the three different temperatures. Strikingly, at 18°C *BG-luc* oscillations in control and *per^I530A^* were of similar amplitude in both LD and DD conditions, with somewhat increased overall expression levels in the mutant (Figure 4A). At 25°C *BG-lu*c oscillations were strongly reduced in the *per^I530A^* mutant background compared to the controls, and this effect was further enhanced at 29°C (Figure 4A). Oscillations in a wild type background were similarly robust at 18°C and 25°C, but amplitude was reduced at 29°C. Next we analyzed the expression of a *tim-luciferase* (*ptim-TIM-LUC*) reporter gene that accurately reflects temporal TIM expression in peripheral clock cells (29, 36). In a wild type background, *ptim-TIM-LUC* oscillations were indistinguishable at 18°C and 25°C, while oscillations were reduced in amplitude at 29°C (Figure S4A), similar to what we observed for *BG-luc* (Figure 4A). This was particularly the case in DD at 29°C, presumably because oscillations of this peripheral clock reporter dampen rapidly in DD even at lower temperatures (Figure S4A) (36). In the *per^I530A^* mutant background, *ptim-TIM-LUC* oscillations were unaffected at 18°C, except for a reduced amplitude in DD (Figure S4A). At 25°C already in LD a strong reduction of TIM amplitude caused by increased trough levels was observed (Figure S4A). In fact, the remaining oscillations are comparable in amplitude to those observed in clock less *per^01^; ptim-TIM-LUC* flies at 25°C, indicating that they are mainly driven by light-dependent TIM degradation during the day (Figure S4B) (37). At 29°C, the mutants’ effects on *ptim-TIM-LUC* expression are further enhanced, leading to largely arrhythmic expression during LD and DD (Figure S4A). In summary, the luciferase reporter experiments confirm that the *per^I530A^* mutant causes a temperature-dependent impairment of clock protein expression, ranging from normal function at 18°C to severe impairments at 29°C.

**Figure 4:**
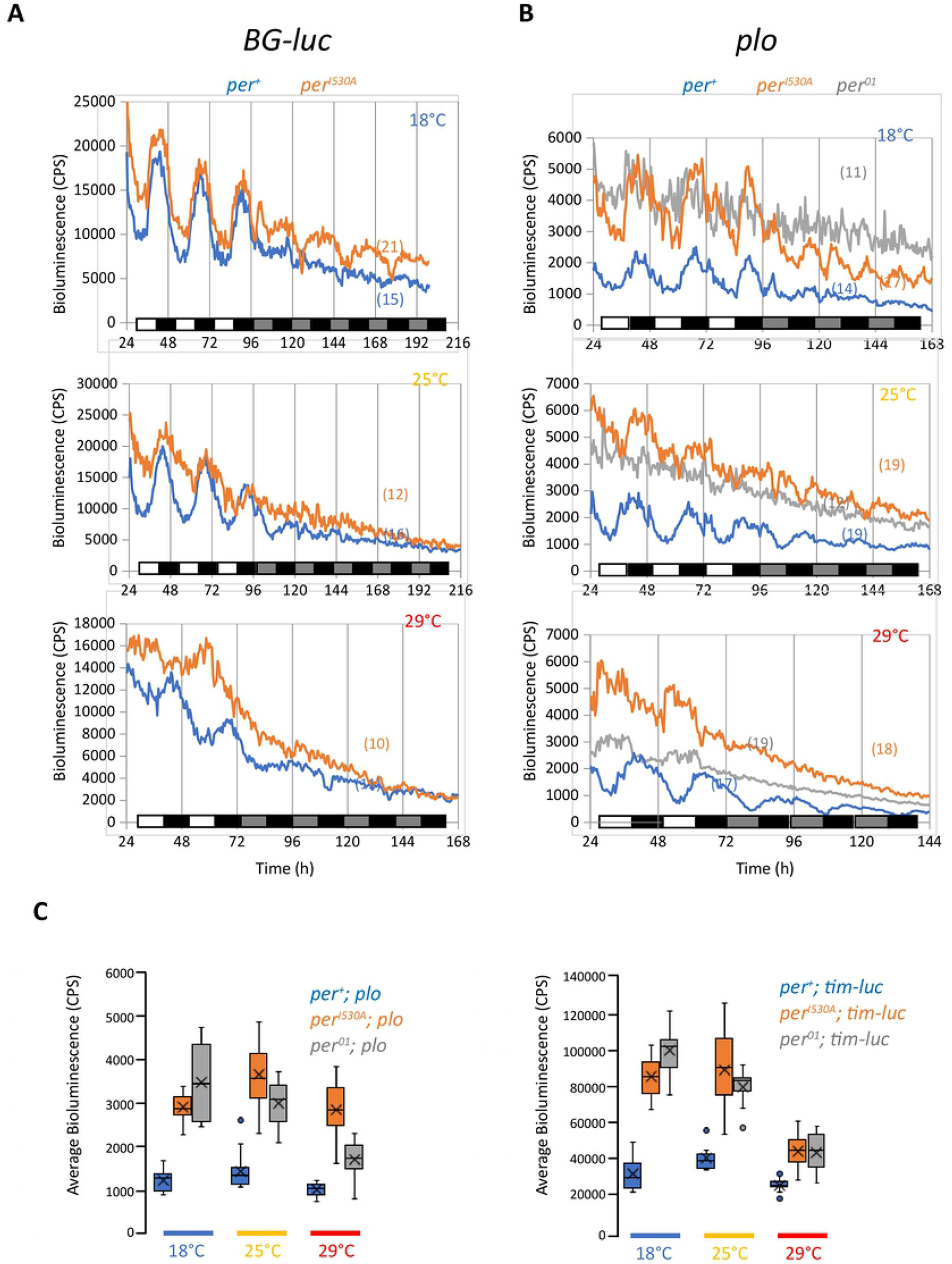
*per^I530A^* causes temperature-dependent dampening of PER-LUC oscillations and impairs repressor function. **(A)** Bioluminescence rhythms of *per^+^* and *per^I530A^* flies expressing the *BG-luc* fusion protein, reporting temporal PER expression. Individual flies were measured at the indicated temperatures for 2 or 3 days in 12 hr : 12 hr LD, followed by 4 or 5 days in DD. White bars indicate lights on, black bars lights off, and grey bars subjective day. N numbers are indicated in parenthesis. **(B)** Bioluminescence rhythms of *per^+^*, *per^01^*, and *per^I530A^* flies expressing the *plo* transgene, reporting temporal *per* transcription and levels. Experiments were performed as in (A). **(C)** Quantification of *plo* (left panel) and *tim-luc* (reporting *tim* transcription, right panel) average bioluminescence levels for the duration of the entire experiments shown in (B) (*plo*) and Figure S4C (*tim-luc*) for the indicated genotypes and temperatures.

### Mutations in the *per^Short^* phosphocluster do not cause temperature-dependent period lengthening

Our Western blot results indicate a temperature-dependent phosphorylation defect caused by *per^I530A^*. Because the isoleucine at position 530 is not a target for phosphorylation by a known PER kinase, we investigated if *per^I530A^* may affect PER phosphorylation by some of the known PER kinases. To this end, we analyzed mutations affecting the PER ‘phosphotimer’ known to set clock speed in *Drosophila* (5). As expected, mutation of the S596 NEMO site (*per^S596A^*) resulted in a drastic period shortening at 25°C (16.5 hr, Table 2), but periods were similarly short at both 18°C (16.9 hr) and 29°C (15.8 hr). Similarly, mutation of a single DBT target site in the *per^Short^* cluster (*per^S585A^*) resulted at short periods at all three temperatures, with an overall 1-hr period shortening at 29°C compared to 18°C (Table 2). Mutating all 4 phosphorylation sites within the *per^Short^* cluster (*per^TS583-596^*) also shortened period at all temperatures with a 1 hr increase in clock speed at 29°C compared to 18°C (Table 2). Finally, mutation of S47A led to the expected period lengthening at 25°C (29.8 hr, Chiu et al 2008), but also at 18°C (29.3 hr) and 29°C (30.3 hr) (Table 2). While these results do not rule out a contribution of the PER phosphotimer in temperature compensation (i.e., mutations within the *per^Short^* cluster show a tendency to increase clock speed with rising temperature), it seems clear that disruption of this timer does not underlie the drastic overcompensation phenotype of *per^I530A^*.

**Table 2:** Free-running behavioral rhythms of *period* mutants with altered phosphorylation sites

| Genotype | Temp (°C) | n | Tau (h ± SEM) | RS ± SEM | % rhythmic |
| --- | --- | --- | --- | --- | --- |
| *per*<sup>S596A</sup> | 18 | 31 | 17.7 ± 0.4 | 4.3 ± 0.2 | 90.3 |
|  | 25 | 32 | 16.9 ± 0.3 | 3.8 ± 0.2 | 100 |
|  | 29 | 27 | 16.0 ± 0.1 | 3.0 ± 0.2 | 88.9 |
| *per*<sup>TS583-596</sup> | 18 | 26 | 17.2 ± 0.1 | 4.7 ± 0.3 | 88.5 |
|  | 25 | 25 | 16.7 ± 0.1 | 4.4 ± 0.2 | 100 |
|  | 29 | 26 | 16.5 ± 0.2 | 3.1 ± 0.2 | 73.1 |
| *per*<sup>S47A</sup> | 18 | 44 | 29.7 ± 0.2 | 3.4 ± 0.2 | 100 |
|  | 25 | 47 | 30.9 ± 0.1 | 4.9 ± 0.2 | 100 |
|  | 29 | 43 | 31.6 ± 0.2 | 2.7 ± 0.3 | 100 |

### period and timeless transcription is enhanced in per^I530A^ mutants

Hyperphosphorylated PER is required for the repression of CLK/CYC mediated *per* and *tim* transcription (4,38,39). To test if the lack of fully hyperphosporylated PER in *per^I530A^* flies at 25°C and 29°C is associated with reduced repressor function we analyzed *per* and *tim* transcription using appropriate *per-luc* and *tim-luc* reporters (40, 41). Bioluminescence emanating from whole flies was recorded as described above in LD and DD conditions at three different temperatures. At 18°C, both control and *per^I530A^* flies showed robust oscillations of *per-luc* expression in LD conditions, which rapidly dampen out in DD (Figure 4B). However, expression levels in the *per^I530A^* mutants were about 2.5 times higher compared to the controls, indicating reduced repressor or enhanced transcriptional activation function of the mutant protein at this temperature (Figure 4B, C). On average, levels of *per* transcription in the *per^I530A^* mutant were somewhat reduced compared to those in a *per^01^* background (Figure 4B, C). At 25°C the amplitude of *per-luc* oscillations in the *per^I530A^* mutants was reduced compared to the controls (Figure 4B). Moreover, expression levels were now higher compared to those in the complete absence of PER, in agreement with an enhanced impairment of PER^I530A^ function at 25°C compared to 18°C. The effect on *per-luc* expression was further enhanced at 29°C, indicated by significantly higher expression levels compared to those in *per^01^* (Figure 4B, C). Furthermore and like in *per^01^*, expression at 29°C was increased during the light phase of the LD cycle in contrast to the clock-dependent *per-luc* peak occurring during the dark phase (Figure 4B) (40). The most likely explanation for this light-dependent peak in *per^01^* and *per^I530A^* flies is the release of *per* transcriptional repression by TIM, caused by light-dependent degradation of TIM, which also occurs in flies without functional PER (see above) (37). We obtained similar effects of the *per^I530A^* mutation using the *tim-luc* transcriptional reporter, although at 29°C there was no difference in *tim* transcription between *per^01^* and *per^I530A^* flies (Figure 4C, S4C). In summary, while at 18°C both genes are rhythmically transcribed in *per^I530A^* mutants, expression becomes progressively disturbed with increasing temperature. In addition, the high levels of *per-luc* and *tim-luc* expression point to an impairment of PER^I530A^ repressor function and/or gain of transcriptional activator function, particularly pronounced at 25°C and 29°C.

### PER:TIM heterodimerization is not grossly altered in *per^I530A^* flies

A possible reason for reduced repressor function of the PER^I530A^ could be the lack or reduction of heterodimerization with TIM. To test if PER:TIM heterodimerization is affected in *per^I530A^* flies, we performed Co-immunoprecipitation (CoIP) experiments using *per^I530A-V5^* and *per^V5^* control flies. Head protein extracts were precipitated with anti-V5 antibodies, and Western blots were subsequently probed with anti-TIM antibodies (Figure 5A, Methods). CoIPs were performed at ZT16, a time when PER and TIM are primarily cytoplasmic, and at ZT22, when both proteins are primarily nuclear (42). To compare dimer formation between mutants and controls in a quantitative manner, we normalized bound TIM to the input and represented it as relative signal intensity. This allows to compare relative binding of TIM to wild type and mutant PER within one temperature. Note that with method, it is not possible to compare absolute levels between temperatures (see Materials and Methods). At 18°C, robust PER:TIM and PER^I530A^:TIM interactions were observed at both time points, potentially somewhat reduced in the presence of the mutant protein, particularly at ZT16 (Figure 5A). Similar results were obtained at 25°C and 29°C, again showing a trend for reduced PER^I530A^:TIM interactions at ZT16 (Figure 5A). Interestingly, at ZT22 this trend was reversed, with higher levels of PER^I530A^:TIM dimers at 25°C and 29°C compared to PER:TIM heterodimers at these temperatures and also compared to PER^I530A^:TIM dimers at 18°C (Figure 5A). The results clearly show that the PER^I530A^ protein is able to form heterodimers with TIM at all temperatures tested, both at ZT16 and ZT22. Faulty PER:TIM heterodimerization is therefore unlikely to explain to observed behavioral and molecular phenotypes of *per^I530A^* flies.

**Figure 5.**
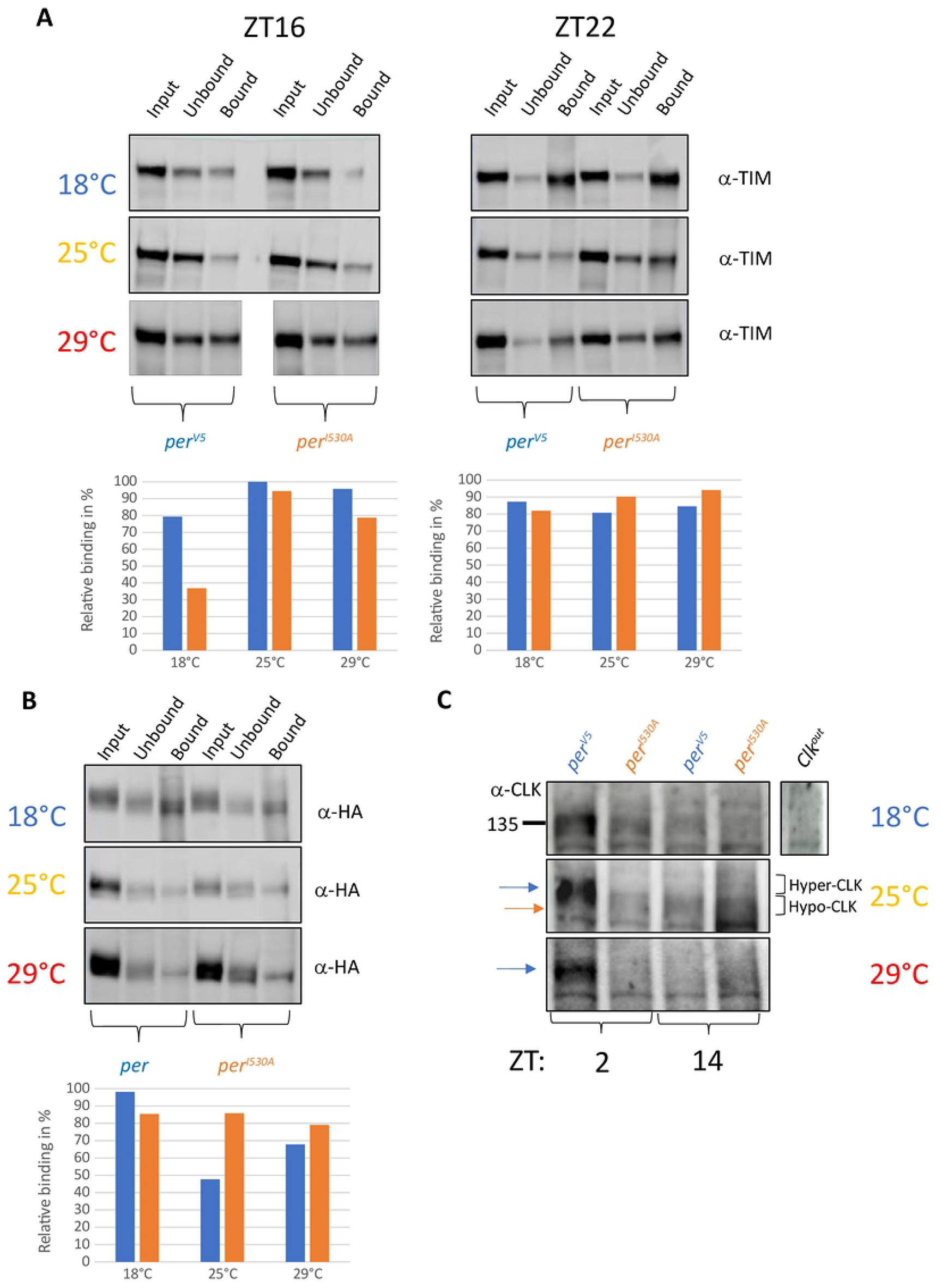
*per^I530A^* does not disrupt PER:TIM and PER:PER dimerization but effects posttranslational modification of CLK. **(A)** PER-TIM interactions in *per^V5^* and *per^I530A-V5^* fly heads at the indicated temperatures. Upper panel: 400μl of fly heads were collected at ZT16 and ZT22 from flies entrained in LD cycles for 3 days and protein extraction was performed as described in Materials and Methods. Protein extracts were loaded onto beads containing anti-V5 antibody. Western blot analysis with anti-TIM antibodies to check TIM binding to PER^V5^ or PER^I530A^, respectively. Input: total protein extract containing PER and TIM. Unbound: supernatant after binding the extracts to anti-V5 coated beads. Bound: proteins eluted from anti-V5 coated beads. Lower panel: bar graphs representing the relative binding levels calculated from three individual experiments. Values are normalized to total amount of protein loaded in the IPs and are expressed as relative signal intensity (highest value = 100%). **(B)** Homodimerisation of PER and PER^I530A^ in fly heads at the indicated temperatures. Upper panel: Flies expressing wild type PER (*per^01^;1-5-1-HA/+;2-2-2-myc/+,* lanes 1-3) or PER^I530A^ (*per^01^;8-6-myc/8-6-myc; 7-3-HA/7-3-HA,* lanes 4-6) encoding *per* transgenes fused to HA and c-myc tags were collected at ZT22 and CoIPs experiments were performed essentially as described above. Beads were incubated with anti-c-MYC antibodies and western blots were developed with anti-HA antibodies. ‘Input’, ‘Unbound’, and ‘Bound’ are as described above. Lower panel: bar graphs representing the relative binding levels calculated from three individual experiments. (C) Western blots comparing CLK expression in heads of *per^V5^* and *per^I530A-V5^* at the indicated temperatures. Flies were entrained in LD cycles for three days and collected at the indicated time points. Extracts from the loss-of-function mutant *Clk^out^* served as a negative control.

### PER:PER homodimerization is not grossly affected in *per^I530A^* flies

In addition to PER:TIM heterodimerization, PER:PER homodimerization is important for PER nuclear entry and PER repressor function (43). We therefore tested if PER^I530A^ is able to form homodimers at different temperatures. For this, we added HA or MYC antigen tags to a full length *per* transgene containing the *per^I530A^* mutation and crossed these transgenes into a *per^01^* background. Wild type versions of these constructs (*per^c-myc^* and *per^HA^*) restore robust behavioral rhythms in *per^01^* flies at 25°C these rhythms are also temperature compensated (43) (Table S1). As expected *per^01^* flies expressing the tagged PER^I530A-MYC^ and PER^I530A-HA^ constructs lengthen their free-running period with increasing temperature, similar to the endogenous *per^I530A^* CRISPR mutants (Table S1). Using Co-IP experiments we have previously shown the abundant existence of PER^HA^:PER^MYC^ dimers in fly heads at 25°C (43). Here we show that PER homodimers also form at 18°C and 29°C (Figure 5B). Next, we compared PER:PER and PER^I530A^:PER^I530A^ dimer formation at the different temperatures in the tagged flies at ZT22 (Figure 5B). The results were very similar to those obtained for PER:TIM heterodimerization: While PER^I530A^:PER^I530A^ homodimers do form at all three temperatures, they appear more abundant at 25°C and 29°C compared to PER:PER homodimers, and this trend is again reversed at 18°C (Figure 5B). Overall, the Co-IP experiments suggest that at ZT22 levels of both PER^I530A^:PER^I530A^ homodimers, as well as PER^I530A^:TIM heterodimers are increased compared to the wild type dimers at 25°C and 29°C. Increased levels of nuclear PER^I530A^ caused by faulty nuclear export offers a potential explanation for this observation.

### Phosphorylation of the transcription factor CLOCK is altered in *per^I530A^* mutants

An important function of PER in the nucleus is to bind to the transcription factor CLK, thereby recruiting kinases that phosphorylate and thereby inactivate CLK, resulting in repression of *per* and *tim* transcription. To see, if the increase *per* and *tim* transcription levels in *per^I530A^* flies are caused by inefficient inactivation of CLK, we compared CLK levels and phosphorylation status at ZT2, a time when CLK is maximally phosphorylated and repressed in wild type flies (reference). Strikingly, in *per^I530A^* flies, CLK phosphorylation was severely affected and mainly hypophosphorylated forms were visible at 25°C and CLK was barely detectable at 29° C (Figure 5C, compare blue and orange arrows in middle panel). Indeed, at ZT2 and 25°C PER^I530A^ migrated indistinguishably from wild type PER at ZT14, a time when CLK is transcriptionally active and hypophosporylated (Figure 5C, orange arrow) (4, 38). In contrast, although CLK levels were reduced in *per^I530A^* mutants at 18°C, CLK migrated similar in wild type and *per^I530A^* mutants, indicating normal phosphorylation and repression of CLK (Figure 5C, black bar). Although PER^I530A^ can form PER:TIM and PER:PER dimers, it appears that its ability to repress CLK (or even binding to CLK) seems to be affected in a temperature dependent manner.

### Subcellular localization of PER^I530A^ is altered at warmer temperatures

If the *per^I530A^* mutation alters nuclear export in a temperature-dependent way, it should be possible to detect differences in PER^I530A^ subcellular localization at different temperatures. To see if this the case, we first looked at wild type PER expression in the small ventral lateral neurons (sLNv), which are pacemaker neurons important for the maintenance of behavioral rhythms and for determining the period length in DD (15). Brains of wild type flies were dissected at two time points: ZT16, when PER is largely hypophosphorylated and cytoplasmic, and ZT22, when PER is largely hyperphosphorylated and nuclear. We found that the levels and subcellular localization (ratio of nuclear versus cytoplasmic signals) of PER were similar at 18°C, and 29°C with expected cytoplasmic and nuclear redistribution between ZT16 and ZT22 (Figure 6A, C, S5). This is in-line with the Western blot results (Figure 3A) and the temperature compensated behavioral period length (Figure 1, Table 1). In contrast, PER^I530A^ showed temperature-dependent variations in nuclear/cytoplasmic distribution. Compared to wild type, nuclear PER^I530A^ levels were drastically reduced at ZT22 and 29°C (Figure S5). Strikingly, at ZT16 and the same high temperature, PER^I530A^ was predominantly nuclear, despite overall lower PER levels (Figure 6, S5). These results indicate that at 29°C, wild type PER enters the nucleus earlier (≤ ZT16) compared to cooler temperatures (∼ZT18) and is then exported again to mitigate premature nuclear accumulation and period shortening at warmer temperatures. We have no explanation for why we observed high PER^I530A^ at 29°C in head extracts, but low levels in the sLNv. One possibility is that in the sLNv premature, nuclear-trapped PER^I530A^, is rapidly degraded, presumably due to aberrant phosphorylation. In contrast, in the photoreceptor cells, which do not contain a self-sustained molecular clock, clearance of premature-nuclear PER^I530A^ may not occur, because it may not be required in peripheral clock cells.

**Figure 6:**
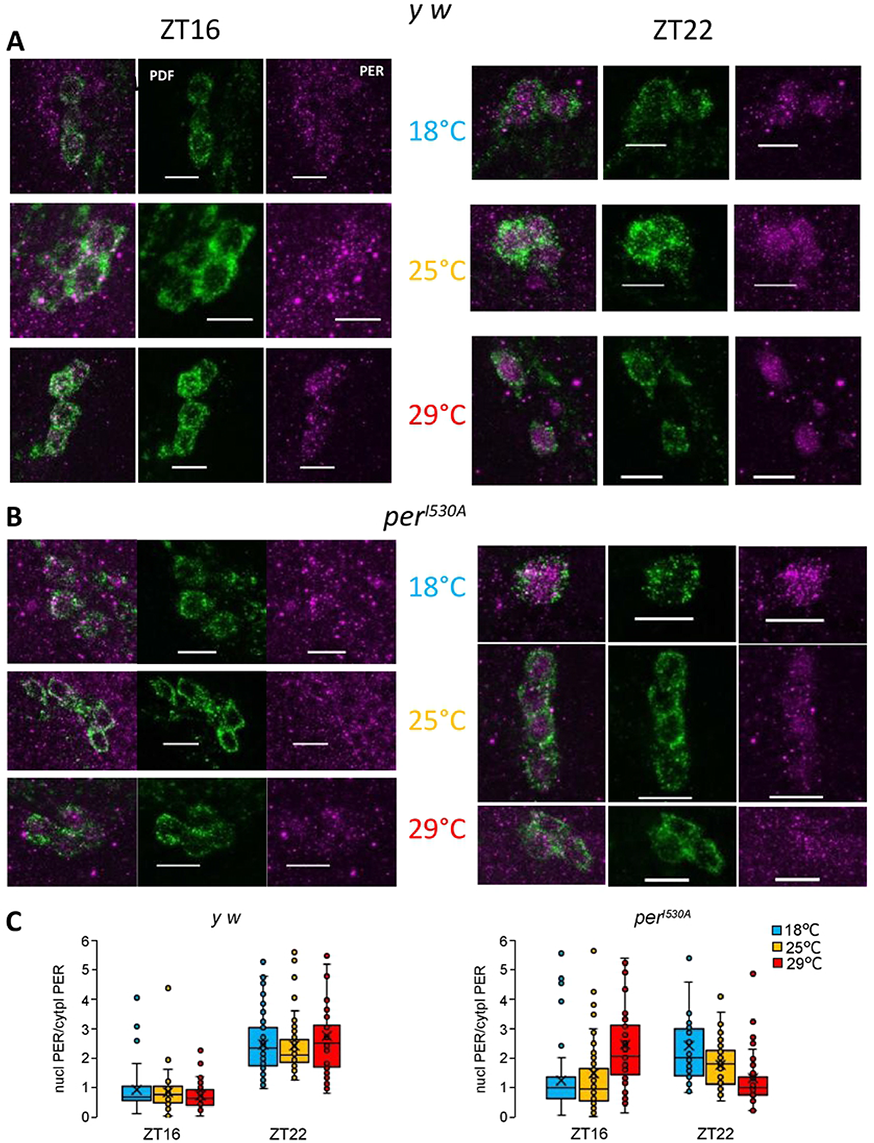
*per^I530A^* modifies subcellular localization of PER in the s-LNv pacemaker neurons in a temperature dependent manner. **(A-B)** Representative images of PER immunostaining in the s-LNv of control flies (A) and *per^I530A^* flies (B) at two time points, ZT16 and ZT22, and 3 temperatures in LD. Cytoplasmic PDF signals were used as a marker to delimitate the nucleus area from the cytoplasmic region. Scale bar: 10 µm. **(C)** Box plots of the fraction of nuclear PER over cytoplasmic PER in individual s-LNv cells in controls (left panel) and *per^I530A^* flies (right panel). Each temperature and time point have been done at least twice. Number of cells for *y w*: 55≤N≤98; Number of cells for *per^I530A^*: 40≤N≤81.

### The nuclear export factor CRM1/Embargoed contributes to temperature compensation

If PER is transported by the general nuclear export machinery mediated by CRM1/Embargoed, knock down of *embargoed (emb*) should also affect temperature compensation. To test this, we used four independent *emb* RNAi lines to knock down *emb* expression in all clock cells (*tim-gal4*), all clock neurons (*Clk856-gal4*), or the subset of the ventral lateral clock neurons (*Pdf-gal4*). To increase the efficiency of RNAi knock-down, *UAS-dicer* was co-expressed with all *RNAi* constructs (44). While control flies containing only the *gal4* or RNAi construct showed stable locomotor activity periods at 18°C, 25°C, and 29°C, this was not the case for lines expressing *emb* RNAi constructs in clock cells and clock neurons (Figure 7A, B, Table 3). Strikingly, and similar to *per^I530A^* mutants, knock down of *emb* led to significantly longer periods at 25°C compared to 18°C. Depending on the driver and the RNAi line the period lengthening varied between 0.7 h and 5 h (average 2.8 h), indicating a massive disturbance of temperature compensation (Figure 7A, B, Table 3). Interestingly, further increasing the temperature to 29°C reversed the effect on clock speed, with periods shortening between 0.5 h and 2.6 hr (average 1.5 h), but not shortening back to the values observed at 18°C (Figure 7 A, B, Table 3). While this effect on temperature compensation is different to the linear increase of period length in *per^I530A^* mutants with temperature, it nevertheless confirms the involvement of nuclear export in temperature compensation. Moreover, it rules out that the observed period lengthening at 25°C is simply caused by increased knock down efficiency due to the temperature dependency of the Gal4 system. The period shortening observed at 29°C most likely indicates that other clock proteins are also subject to temperature dependent nuclear export. To directly test the interaction between specific knockdown of nuclear export and the *per^I530A^* mutation, we analyzed the behavior of flies expressing *emb-RNAi* in the *per^I530A^* background (Figure 7C, Table 4). Because we could not co-express *UAS-dicer* as in the *RNAi* experiments described above (both *UAS-dicer* and *per^I530A^* are located on the *X-* chromosome), we compared the period length of the double mutants to those of *per^I530A^* single mutants carrying the *gal4* only. Using *Pdf-gal4* to drive two different *emb-RNAi* constructs in *per^I530A^* mutant background resulted in a significant blunting of the temperature-dependent period increase observed in *per^I530A^* single mutants (Figure 7C, Table 4). Moreover, at 25°C expression of *emb-RNAi BL34021* resulted on 100 % arrhythmic flies, which is not observed in *per^I530A^* single mutants or *Pdf > emb-RNAi* flies in a *per^+^* background (Tables 1, 3, 4). Using *tim-gal4* to drive *emb-RNAi BL31353* in a *per^I530A^* mutant background also blunted temperature-dependent period increase compared to *per^I530A^*, *tim-gal4* flies (Figure 7C, Table 4). Taken together, these results indicate an interaction between nuclear export and the *per^I530A^* mutant and suggest that in addition to PER, nuclear export of other clock proteins is regulated in a temperature-dependent manner (see Discussion).

**Figure 7:**
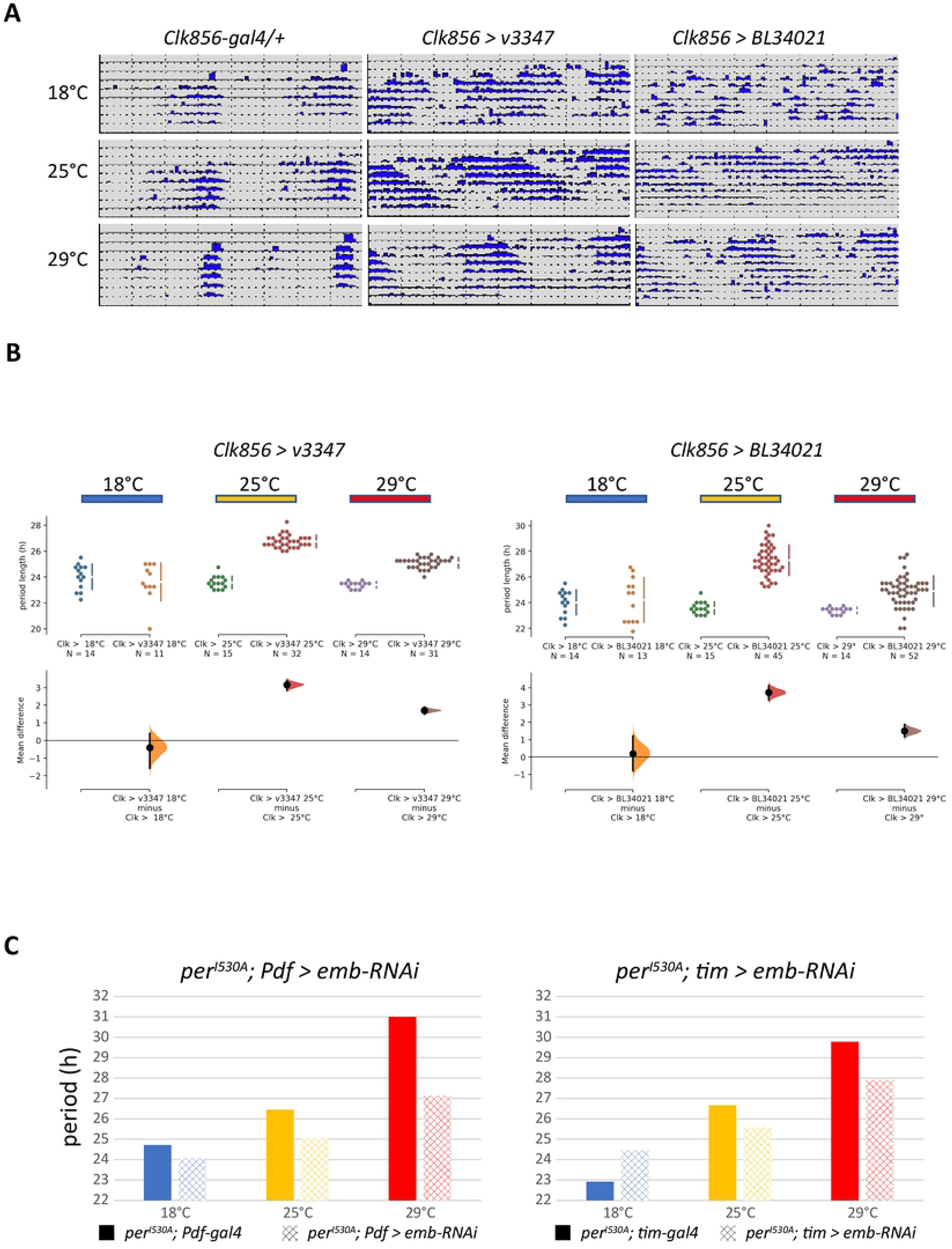
The nuclear export factor CRM1/EMBARGOED affects temperature compensation. **(A)** Effect of downregulating *embargoed* (*emb*) in clock neurons on free-running locomotor activity analyzed at different temperatures. *UAS-dicer Clk856-gal4* was used to downregulate *emb* using two different RNAi lines (BL34021 and v 3347). Flies were kept for 3-4 days in LD followed by an additional week in DD at the constant temperatures indicated on the left side of each panel. Individual actograms of the LD and DD part of the experiments are shown at the three temperatures. **(B)** Quantification of period length and differences between control (*UAS-dicer; Clk856-gal4/+*) and *emb* knockdown flies (*UAS-dicer; Clk856-gal4/+; v3347/+*) and *UAS-dicer; Clk856-gal4/BL34021*) using ES (see legend to Figure 1C and Materials and Methods for details). For additional *gal4* driver and *emb-RNAi* lines see Table 3. **(C)** Free running period length of flies with *emb* downregulation in the background of the *per^I530A^* mutation. Behavior was analyzed as in (A) and average periods compared between controls (*per^I530A^*; *Pdf-gal4/+* or *per^I530A^*; *tim-gal4/+*) and *per^I530A^ emb-RNAi* knockdown flies (*per^I530A^*; *Pdf-gal4/+; BL31353/+* or *per^I530A^*; *tim-gal4/+; BL31353/+*) at the indicated temperatures. Error bars indicate SEM. For additional *emb-RNAi* lines see Table 4.

**Table 3:**
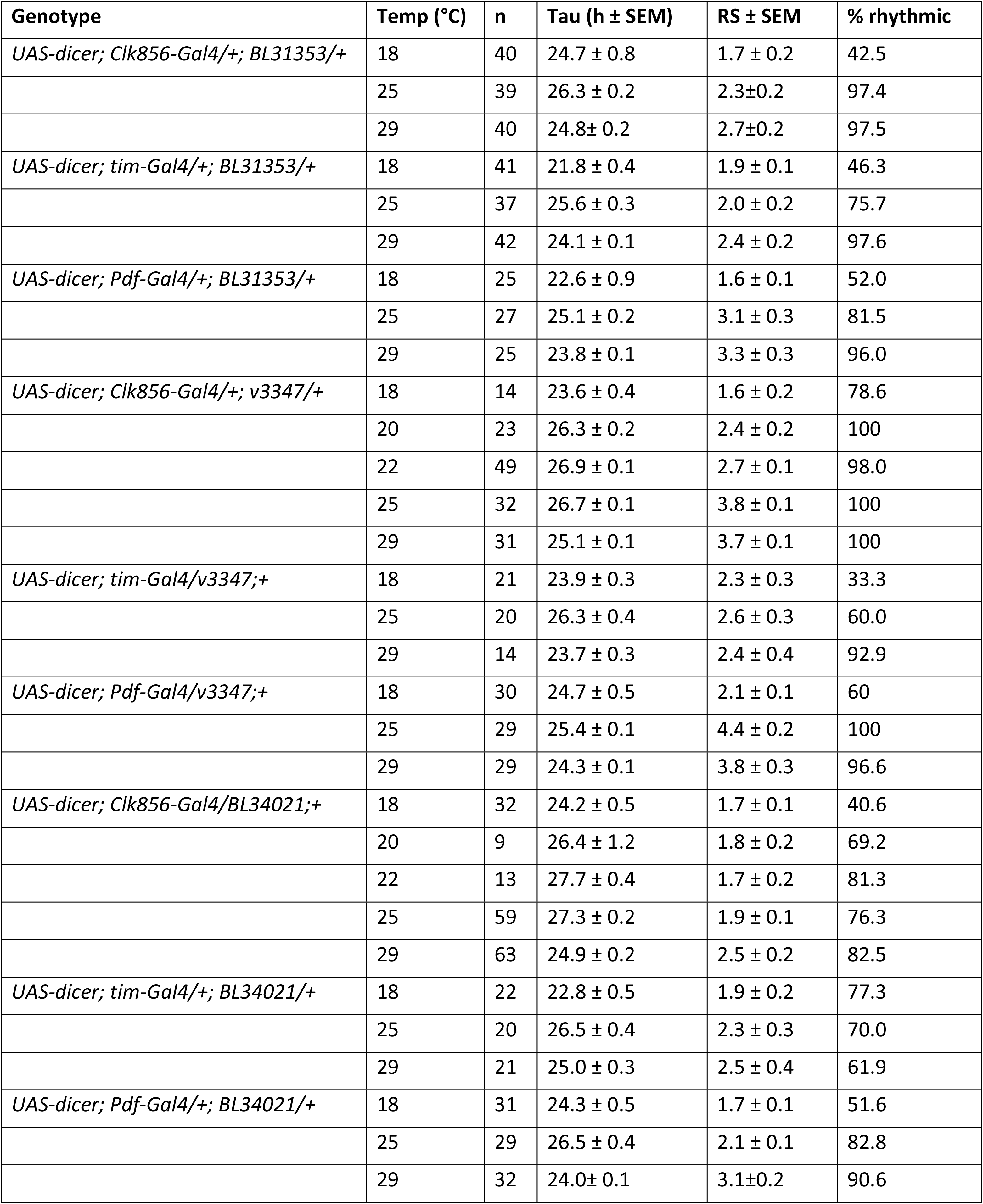
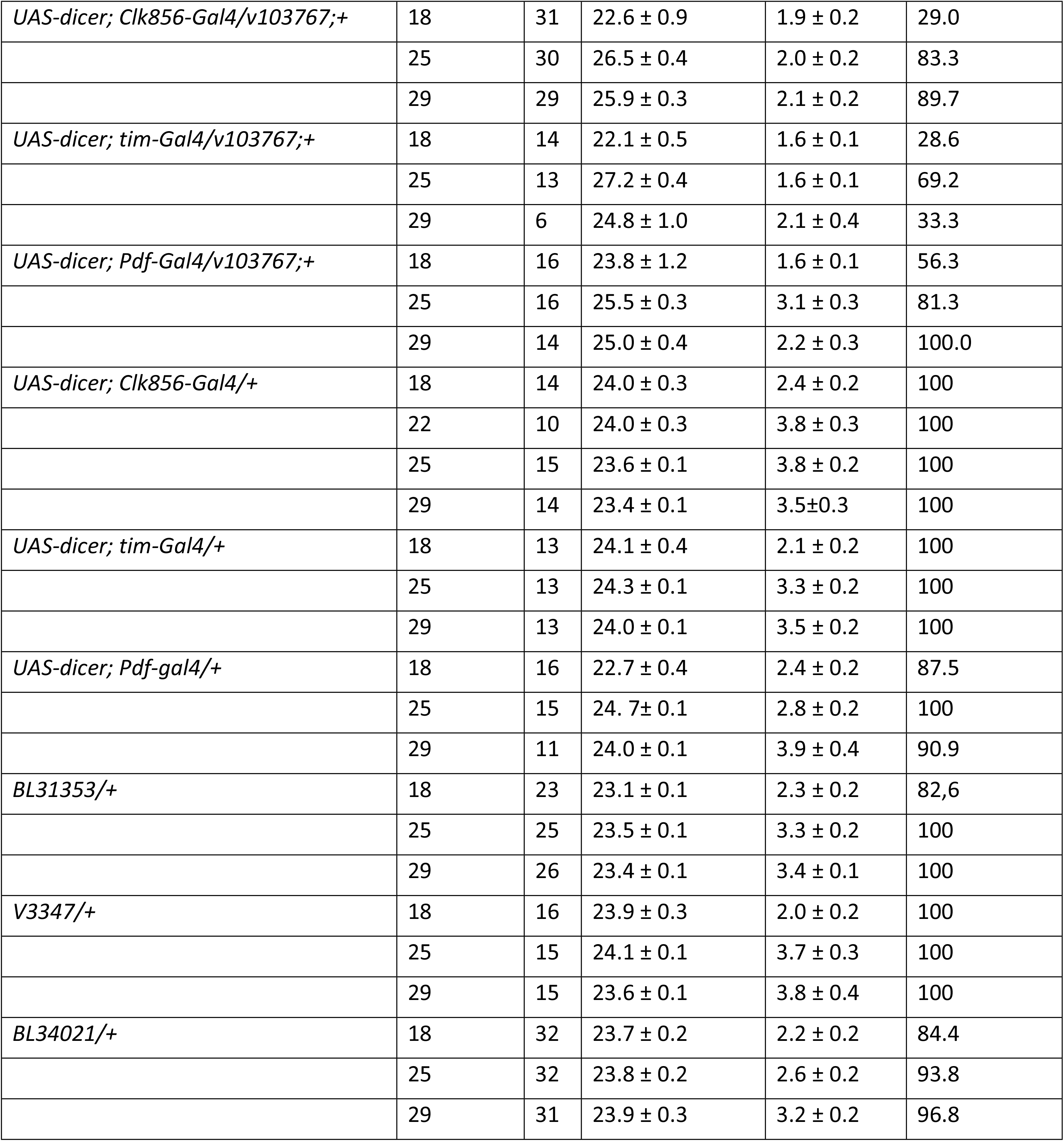
Free-running behavioral rhythms of *embargoed* RNAi lines and controls

**Table 4:**
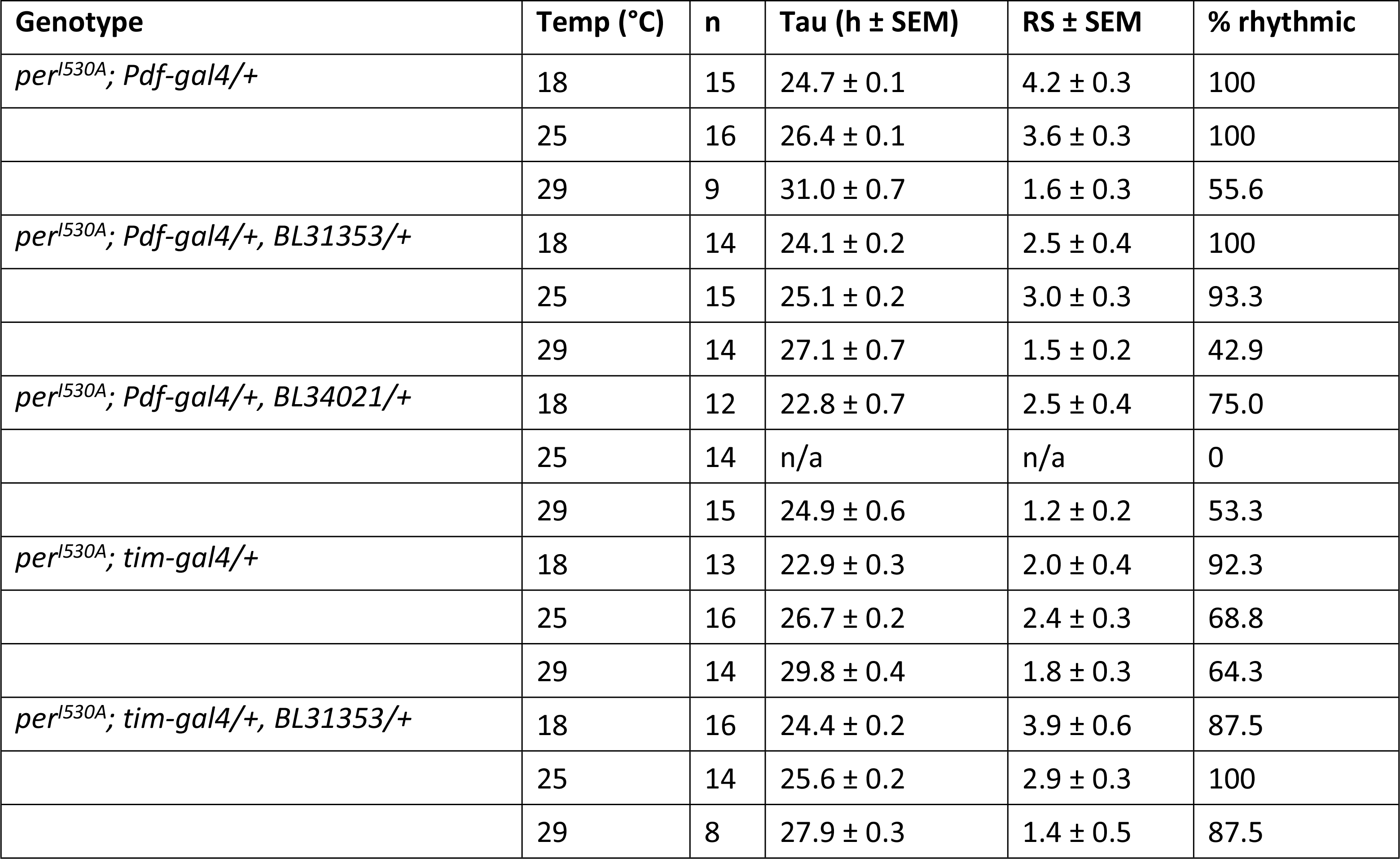
Free-running behavioral rhythms of *per^I530A^ emb-*RNAi double mutants and controls

## Discussion

Circadian clocks are able to compensate for the acceleration of biological reactions associated with an increase in temperature. An impairment of temperature compensation should therefore result in a period shortening with increasing temperature. The *per^I530A^* mutation has the opposite effect, indicating that this mutation leads to an overcompensation phenotype. Nevertheless, the molecular defects associated with *per^I530A^* could shed light on the mechanism underlying temperature compensation, and indeed our findings indicate that nuclear export plays an important role in this process. First, nuclear levels of both PER^I530A^:TIM heterodimers and PER^I530A^:PER^I530A^ homodimer levels appear increased in *per^I530A^* mutants compared to controls, which could be explained by reduced nuclear export efficiency of the mutant protein (Figure 5A, B). Second, subcellular localization of PER^I530A^ revealed a temperature-dependent atypical nuclear enrichment at ZT16 at 29°C, in agreement with the idea that wild type PER is exported from the nucleus at this temperature to mitigate early nuclear accumulation. Finally, knockdown of the nuclear export factor CRM1 effected temperature compensation in opposite ways. In the range from 18°C to 25°C *emb-RNAi* expression resulted in a significant lengthening of period with the same magnitude (about 3-4 h) as observed in *per^I530A^* flies (Figure 7B, Table 3). In contrast, further increase of temperature to 29°C lead to a significant period-shortening by ∼1.5 h. These results unequivocally show that nuclear export is involved in the temperature compensation mechanism. Moreover, they indicate that in addition to PER, other clock proteins are subject to nuclear export. At low temperatures (18°C) nuclear export does not seem to play a role in adjusting period length, as both *per^I530A^* and *emb-RNAi* flies show periods in the normal 24 h range. At 25°C, both *per^I530A^* and *emb-RNAi* flies show essentially the same period-lengthening phenotype, indicating that in wild type flies, PER needs to be exported out of the nucleus in order to maintain a normal period length. At 29°C *per^I530A^* flies further increase their period length, indicating the necessity for continued temperature compensation of PER nuclear export above 25°C. In contrast, *emb-RNAi* flies shorten their period length from 25°C to 29°C by more than one hour. This suggests that at least one other clock protein needs to be exported at temperatures above 25°C to avoid speeding up the clock at temperatures above 25°C. A good candidate would be the unknown CLK-kinase, required to shut down CLK transcriptional activity (4, 39). Increased kinase activity with rising temperature would be counteracted by increased nuclear export of the kinase. This idea is also consistent with the phenotype of flies with knocked down *emb* expression in the *per^I530A^* mutant background: at 29°C this resulted in reduction of the *per^I530A^*-induced period lengthening by 1-2 h, while there was little effect at 25°C (Figure 7C, Table 4). Since mutations affecting TIM NES function also show temperature dependent period lengthening from 18°C to 29°C, it is unlikely that blocking TIM nuclear export is responsible for the period shortening observed in *emb-RNAi* flies above 25°C. We therefore propose that PER and TIM (presumably as PER:TIM heterodimer and PER:PER homodimer) are both subject to increased nuclear export with increasing temperature.

### Molecular consequences of nuclear PER^I530A^ retention at higher temperatures

The temperature-dependent phosphorylation defect of PER^I530A^ (Figure 3B) combined with its reduced repressor activity (Figure 4B, C) point to an impairment of this protein to recruit kinases to the CLK transcription factor, which are important for inactivating this transcription factor (4, 39). Based on the striking phosphorylation defect of PER^I530A^ at 25°C and 29°C (Figure 3B), we propose that the mutant protein is retained in the nucleus and ‘protected’ from kinases that would normally phosphorylate PER in the cytoplasm to promote nuclear entry (45–48). At 18°C, where nuclear export is less important, PER^I530A^ would be normally phosphorylated and imported into to the nucleus. If true, this implies that wild type PER proteins are shuttled in and out of the nucleus at warmer temperatures to prevent early nuclear accumulation and speeding up of the molecular feedback loop with increasing temperature. Overall a relatively simple model emerges in which temperature-dependent enzymatic events (e.g. kinase activities) leading to earlier accumulation of PER and TIM in the nucleus and presumably to premature inactivation of CLK at warm temperatures (> 25°C), are counteracted by increased nuclear export. Although possibly too simplistic, this model is attractive, because it concedes that all enzymes and other factors involved naturally increase their reaction rate and activity with increasing temperature.

### Temperature compensation: Network versus cell autonomous property

The *Drosophila* circadian clock network driving behavioral activity rhythms consists of several coupled neuronal groups (14). Following Pittendrigh’s proposition, it is therefore conceivable that in order to compensate temperature changes some of these groups speed up with temperature, while others slowdown (10). While such a scenario is possible, available evidence points to temperature compensation being a cell autonomous property, affecting distinct steps in clock regulation as suggested by others (18). Using the novel LABL technique, we were able to monitor *period* promoter activity within specific subsets of the clock network and found that all groups are properly temperature compensated. Importantly, this included the DN1 neurons, which on their own are not able to sustain behavioral rhythmicity in DD, as well as peripheral clock cells (Figure 2A, B). Although it can be argued that signals were recorded from specific neuronal subsets, these subsets within the context of the entire network, which could of course influence the oscillations within one group. Nevertheless, even in the background of a *Pdf^01^* mutation, which largely prevents network coupling due to the lack of the PDF neuropeptide (17), temperature compensated clock gene expression was observed in dorsal subsets of the clock neurons (29). However, the same study revealed that circadian clocks in isolated fly tissues (in halteres and antennae) are over compensated, pointing to differences between neuronal and non-neuronal circadian clocks (29). Taken together, the currently available data support a model of cell autonomous temperature compensation, at least in central circadian clock neurons.

## Conclusions

The *per^I530A^* NES mutation equally effects temperature compensation on a behavioral and molecular level. Combined with the temperature compensation defects observed after knockdown of the nuclear export factor CRM1, this strongly implicates nuclear export of PER and other clock proteins as important part of the underlying molecular mechanism. Nuclear retention of PER^I530A^ leads to increased CLK-mediated transcriptional activity, most likely caused by a temperature-dependent PER^I530A^ phosophorylation defect. Our results support a model in which premature nuclear accumulation and CLK-inactivation at warmer temperatures is counteracted by increased nuclear export of clock factors, resulting a temperature compensated circadian clock. Future work will reveal, which other clock factors show temperature dependent control of subcellular localization.

## Acknowledgements

We thank Patrick Emery for discussions and for sharing unpublished results. We thank Isaac Edery and Joanna Chiu for TIM and CLK antibodies, Paul Hardin for *Clk^out^* flies, Paolo Sassone-Corsi for the pAc5.1-I530A clone, as well as Mechthild Rosing for technical support. This work was supported by a grant from the Deutsche Forschungsgemeinschaft given to RS (STA 421/7-1) and a grant from NSERC (RGPIN-2019-06101) to D.T.

## Materials and Methods

### Flies

*Pdf*-*gal4* (49), *DvPdf*-*gal4* (50), *R18H11-gal4* (51), *tim-UAS-gal4* (30), *tim-gal4:27* (52), *Clk856-gal4* (53), *Clk^out^* (38), *per^01^* (19)*, per^01^;1-5-1-HA; 2-2-2-myc* (43), *plo* (40), *BG-luc* (31), *tim-luc* (54), and *tim-TIM-LUC* (36) have been described. *UAS-dicer* (inserted on the *X-* chromosome)*, nos-Cas9,* and *emb-RNAi* lines *BL31353* and *BL34021* were ordered from the Bloomington Stock Center or from the VDRC (*v3347* and *v103767*). All other fly lines were generated as part of this study.

### Generation of *per^I530^* and V5-tagged versions of wild *per* and *per^I530^*

#### gRNA design and cloning

Target gRNA sites were selected so that Cas9 mediated cleavage was directed to a target locus of 100bp upstream and downstream of the per I530A mutation. To avoid off target cleavage optimal target sites were identified using CRISPR target finder (http://flycrispr.molbio.wisc.edu/tools). 2 gRNA targets were chosen that are close to the target locus. Complementary target sites oligos also contained a 5’ guanine for transcription from the U6 promoter and a 3 bp overhang compatible to BbsI sites. Oligos were annealed using standard primer annealing reactions and cloned into BbsI linearized pCFD3 plasmid (55) via T4 DNA ligation.

#### Donor plasmid construction

Donor plasmids that contain the desired *per* mutations and all elements necessary for homologous recombination were constructed in three subsequent cloning steps. In each round of cloning the 1.5 kb 5’ homology arm and the 1.5 kb 3’ homology arm were individually PCR amplified from *y w* flies using outside primers HAXbaIF and HAHindIIIR in combination with respective internal primers. Outside primers HAXbaIF and HAHindIIIR contain a 15 bp overhang for In-fusion cloning that is homolog to linearized vector ends. Inside primers have a 5’ 15-20 bp extensions that are complementary to each other in addition to one defined mutation for each round of cloning. In the initial round of cloning Pam site mutations were introduced to avoid unwanted Cas9 cleavage within the donor plasmid. The two fragments (5’ homology arm and 3’ homology arm) were assembled into plasmid pBS-KS-attB1-2-PT-SA-SD-0-2xTY1-V5 (Addgene) that was linearized with XbaI and HindIII using In Fusion cloning. In a second round of cloning the homology arms were amplified again using the pBS donor plasmid from the previous round as a template. Outside primers were as described above while the inside primers introduced either a silent HindIII site that can be used to screen for transformants or the V5 tag. In fusion cloning was used to assemble the fragments as described above. The resulting plasmid was then used in a final round of PCR to introduce the per I530A mutation. See table_ for a detailed list of all primers.

### Embryo injection and screening for transformants

Donor plasmids containing the desired mutation along with gRNA plasmids were verified by sequence analysis and scaled up for injections using Qiagen plasmid midiprep. 6 µg of each plasmid were precipitated and eluted in injection buffer. gRNA construct and donor plasmids were mixed prior to injection and the mix was injected into freshly laid embryos of nos-Cas9 flies that were crossed to *y w* flies (55). Surviving adults were backcrossed in batch crosses to *FM7*; +; + flies to balance *X*-chromosome modifications with *FM7*. Individual male and female flies from this cross were crossed again to *FM7*; +; + flies. After letting the females lay eggs for 3-5 days adult transformant flies were used for molecular screening. Two different strategies were used to screen for transformants. In general, a total of 95 flies for each mutation were screened using either PCR or PCR in combination with restriction digests. For *per* constructs containing the V5 tag a simple PCR approach was used: a ∼260 bp target locus containing the desired mutations was amplified by PCR using genomic DNA from individual flies. Resulting products were analyzed on agarose gels. PCR products of samples that showed an increase of size due to the V5 tag being present were then sequenced to verify the presence of the desired mutations and the V5 tag. For *per* constructs without the V5 tag a ∼800 bp target locus was amplified by PCR using genomic DNA from individual flies. 1ul of HindIII was then added to half of the PCR reaction and incubated for 2 h at 37°C. Resulting products were analyzed on agarose gels. The remaining PCR product of samples that showed digested products of the correct size were then sequenced to verify the presence of the desired mutations.

### Generation of *per^I530A^* versions tagged with c-Myc and HA epitopes

In order to perform the PER:PER homodimer assays it was necessary to generate transgenic flies expressing differentially tagged version of *per^I530A^*. Constructs encoding wild-type C-Myc and HA tagged version of PER have been described previously (43) and contain the full length tagged *per* cDNA, expressed under the control ∼1.3 kb 5’-upstream and 2.1. kb 3’ downstream genomic regulatory sequences. To generate constructs encoding C-Myc and HA tagged PER^I530A^, a 1.1 kb SanDI/BamHI fragment of the two wild type constructs was replaced with the same fragment containing the I530A mutation (gift from Paolo Sassone-Corsi) exactly as described in (43) and the mutation in the final construct was verified by sequencing. Transgenic flies were generated in a *y w* background using classical transposase mediated germline transformation (43). Transgenic flies containing *per^I530A-myc^* on chromosome *2* and *per^I530A-HA^* on chromosome *3* were combined and introduced into *per^01^* genetic background using standard genetic crosses.

Table: Primer list

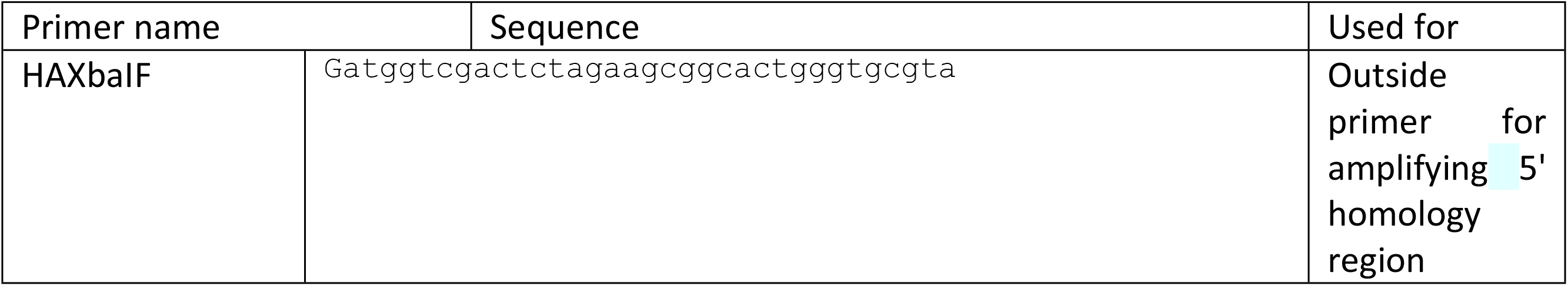
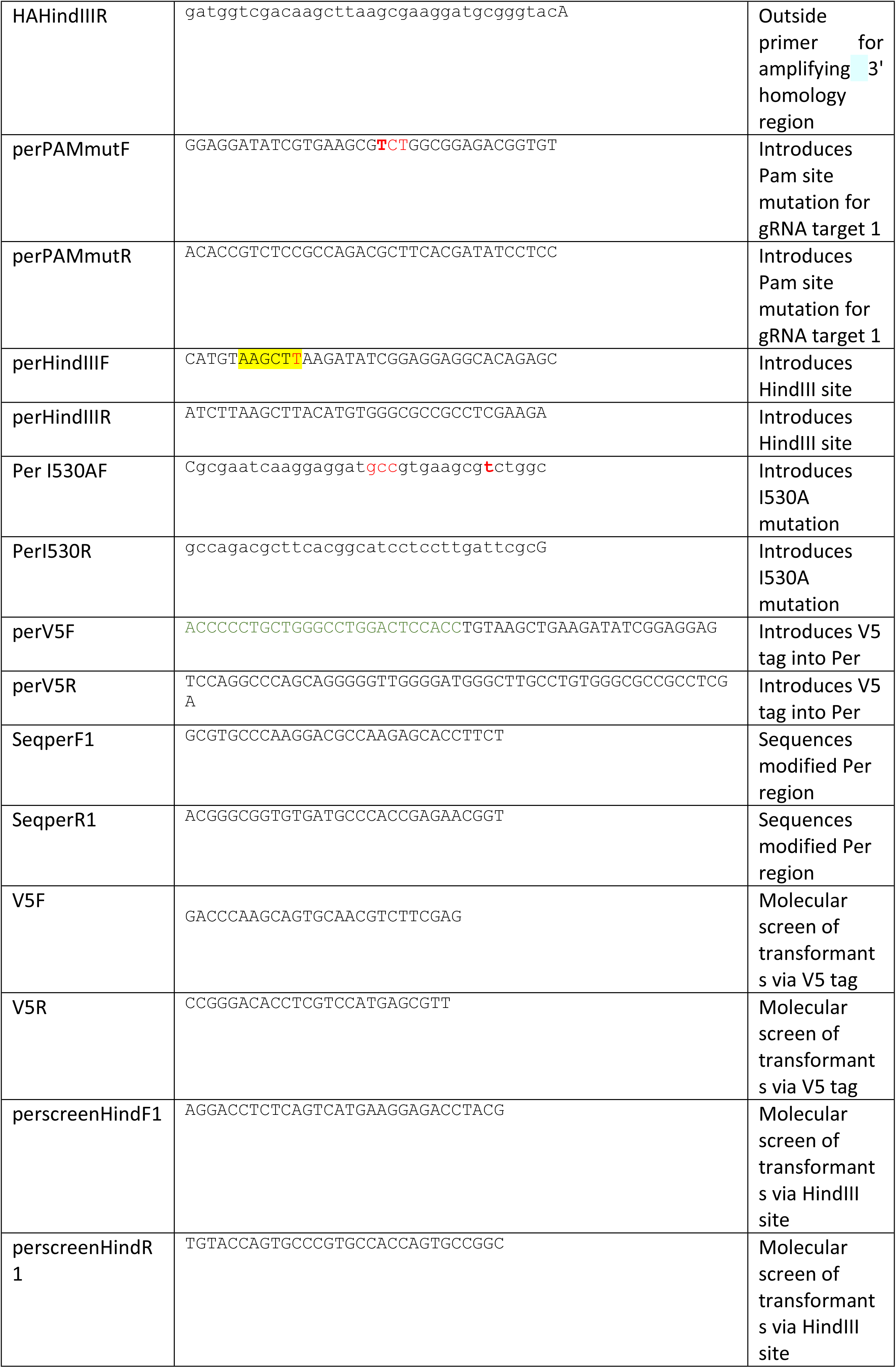
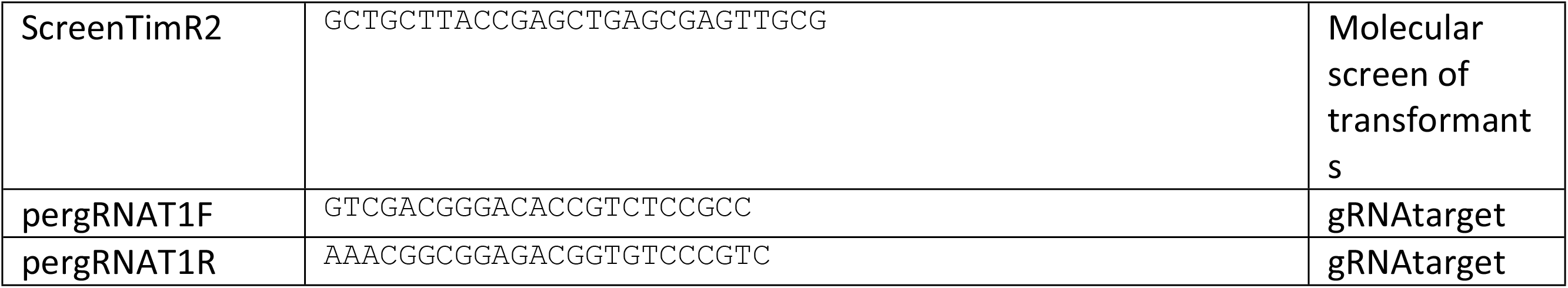

### Behavior

Two to four days old males were loaded into glass tubes containing 5% sucrose in 2% agar and loaded into the DAM2 TriKinetics system (Waltham, MA). Flies were exposed to LD for 3 days, followed by 5-7 days in DD to assess their free running periods at constant temperatures of 18°C, 25°C or 29°C. Period length and their significance (RS values) were determined using autocorrelation and Chi-square periodogram analysis functions of the fly tool box implemented in MATLAB (MathWorks). Period values with associated RS values ≥ 1.5 were considered rhythmic (56). For statistical analysis shown in Figures 1C and 7B, estimation statistics has been used. This approach gives a more informative way to analyze and interpret results (57). It focuses on the effect size, as opposed to significance testing. While significance testing (p-values) focus on the acceptance or rejection of the null hypothesis, estimation stats focus on the magnitude of the effect size (*i.e*. mean difference) and its precision (57). Data were analyzed using DABEST (57), using the website available under https://www.estimationstats.com/#/ as described (29).

### LABL analysis and classical bioluminescence assays

#### LABL

Detailed methods are available in Johnstone et al., 2021 of this issue. Expression of Flipase using the indicated *gal4* drivers allows anatomical restriction of the firefly luciferase reporter (Luc2, Promega). Luminescence of flies were measured using the LumiCycle 32 Color (Actimetrics). D-luciferin potassium salt (Gold Biotechnology) was mixed with standard fly food to a final concentration of 15 mM in Drosophila culture plates (Actimetrics). Analysis software was used to normalize the exponential decay, data were exported into .csv files (Actimetrics) and locally written python code was used to organize luminescence data into 30-minute bins (LABLv9.py; www.top-lab.org/downloads), and to quantify periods of oscillations using a Morlet wavelet fit (waveletsv4.py; www.top-lab.org/downloads). Data were plotted using Graphpad Prism 9.

#### Classic Bioluminescence assays

Luciferase expression of individual flies carrying various *period-luciferase* and *timeless-luciferase* reporter genes was measured as described previously (41). 3-4 day old males were loaded in 96-well microtiter plates containing 100 µl of 5% sucrose, 1% agar and 15 mM luciferin. Bioluminescence was detected with a TopCount Multiplate Reader (Perkin Elmer) for serval days during LD and DD conditions at the designated constant temperature. Bioluminescence was measured twice per hour and data were plotted using BRASS ((Version 2.1.3) (58)).

### Western Blot

V5-tagged flies were kept in LD cycles for 3 days at constant temperatures of 18°C, 25°C or 29°C. On the fourth day, flies were collected at the respective time points and frozen in liquid nitrogen. For DD experiments flies were kept in constant darkness for an additional day before collecting sample collection at the respective times. Protein extractions of adult fly heads were performed as previously described (43). For the V5 and TIM blots ∼30 fly heads were homogenized with 50 µl of Extraction buffer (20 mM HEPES [pH 7.5], 100 mM KCl, 5% glycerin, 10 mM EDTA, 0.1% Triton-X 100, 5mM DTT, 1x Complete Protease Inhibitor and 4mM Pefabloc. For anti-Clk blots RBS buffer (20 mM HEPES [pH 7.5], 50 mM KCl, 10% glycerin, 2 mM EDTA, 1% Triton-X 100, 0,4% NP40, 1mM DTT, 1x Complete Protease Inhibitor and 4 mM Pefabloc) was used for extraction. After homogenization samples were sonicated in 3-5 pulses. Protein concentration was measured using Coomassie Plus Protein Assay Reagent (Thermo scientific) and equal amounts of protein were loaded onto 6% SDS gels for anti-V5 and anti-TIM blots or 8% SDS gels for anti-CLK blots respectively. For anti-V5 and anti-TIM blots proteins were transferred onto PVDF membranes while for anti-CLK blots proteins were transferred onto Nitrocellulose membranes using a Semi-dry electro blotting unit at 240 mA for 40 min. After blotting, membranes were blocked with 5% milk in TBS-T at room temperature for 1 h. First antibodies (5% nonfat milk/TBS-T) were incubated at 4° C overnight, followed by secondary antibody (5% nonfat milk/TBS-T) incubation at room temperature for 1,5 h. Dilution of first antibodies were 1:5000 for mouse anti-V5 (Thermo Fisher), 1:2000 for guinea pig anti-TIM (59), 1:2000 for mouse anti HA-tag (6E2 Cell signaling), and 1:2000 for anti-CLK (3). The dilutions of HRP-conjugated secondary antibodies were 1:2000 for goat-anti-guinea pig IgG and 1:2000 for anti-mouse IgG (Cell Signaling). Signal Fire ECL substrates (Cell Signaling) were used to develop the membranes. To calculate relative expression levels the individual bands were quantified using Image J. The background was subtracted for each individual lane and the amount of protein was expressed as relative signal intensity by dividing each value by the highest value for each blot (highest value = 100%) within one blot. Average values were then calculated from the three individual blots and were blotted as line graphs (± SEM).

### Immunoprecipitations

CoIPs were performed as previously described (43). Briefly, adult flies from V5-tagged strains (*per^V5^* and *per^I530A-V5^*) were entrained to a 12-h:12-h LD cycle for 3 days at constant temperatures of 18°C, 25°C or 29°C. At ZT 16 or ZT20 6 ml of flies were collected and frozen in liquid nitrogen. Fly heads were separated by repeated vortexing/cooling in liquid nitrogen and 400 µl of fly heads were then separated using a 0.45 mm/0.14 mm metal mesh. Heads were then homogenized in 400 µl of Extraction Buffer (20 mM Hepes [pH 7.5], 100 mM KCl, 1 mM DTT, 5% glycerol, 0.05% NP40, 1x Complete Protease Inhibitor and 4 mM Pefabloc.

For the input sample, 20 µl are aliquoted and added with 20 µl of 2x SDS-sample buffer. The remaining head extract was incubated with Anti-V5 Agarose Affinity Gel (Sigma) ON at 4°C. Beads were spun down by centrifugation, 20 µl of supernatant was aliquoted and served as the supernatant (unbound) control. The beads containing the bound fraction were washed three times with Extraction Buffer and subsequently resuspended in buffer for further analysis. Co-IPs done to ckeck for homodimers were done the same way except that agarose beads (Protein G Sepharose™ 4 Fast Flow by GE Healthcare) were used that were incubated with mouse anti-myc antibody (9B11, Cell Signaling) for 1h at 4°C prior to incubating it with the head extracts. The coated beads were then washed with 1 ml of Extraction Buffer, resuspended in 40 µL Extraction Buffer/sample and loaded with homogenized head extracts. Individual bands were quantified using Image J and the background was subtracted for each individual lane. Bound protein was normalized to the input and presented as relative signal intensity by dividing each value by the highest value for each blot (highest value = 100%). Average values were then calculated from three individual blots and were blotted as bar graphs (± SEM).

### Immunostainings

The protocol used was the same as previously described (60). Flies were entrained in LD for 6 days. Flies were collected in the dark and stayed in the dark until dissections. Brains were dissected in PBST 0.1% and fixed for 20 min at room temperature in PFA 4%. After 3 washes brains were blocked for one hour at room temperature in PBST 0.1% + 5% goat serum. Primary antibodies were incubated for 48 h (in PBST 0.1% + 5% goat serum) at 4°C, while secondary incubation was done overnight at 4°C. Brains were mounted using Vectashield. Monoclonal anti-PDF (DSHB) was used at 1/1000, and pre-absorbed Rabbit anti-PER (61) was used at 1/15000. Secondary antibodies used: goat cross absorbed anti-mice 488 ++ 1/2000 (Invitrogen), goat anti-rabbit 555 1/2000 (Invitrogen). Brains were imaged with a Leica TCS SP8 confocal microscope with a 63x objective. Average intensity was measured using ImageJ and quantification was normalized to the background: (signal-background)/background 50.

---

**Figure S1.**
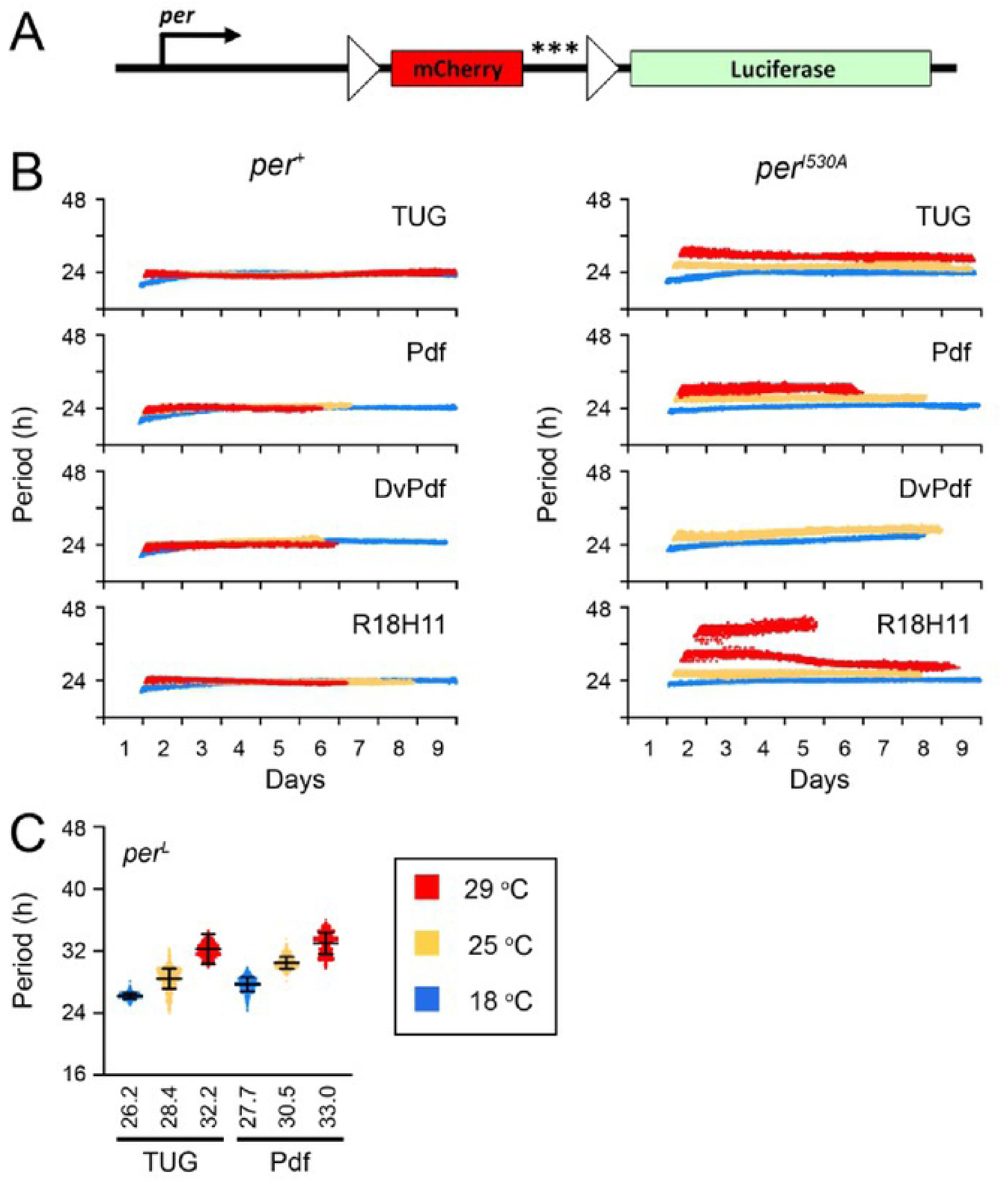
Period of clock oscillations over time, and average clock periods of *per*^L^: (**A**) The architecture of the LABL reporter construct. Details of the construction of this reporter are reported in Johnstone et al., 2021 of this issue. The *period* promoter is fused to mCherry with three stop codons (asterisks) and subsequently to Luciferase. The mCherry gene is flanked by FRT recombination sites (triangles), which permit Gal4-mediated flippase excision of the gene, allowing Luciferase expression by the *per* promoter. (**B**) Changes of period of clock oscillations in wild type flies, over time. LABL is activated using the indicated Gal4 driver in wild type (*per*^+^) and mutant (*per^I530A^*) flies, and changes in period of oscillation calculated by Morlett-wavelet-fitting over time. Colors represent different temperatures: 18 °C (blue), 25 °C (yellow) or 29°C (red). Two replicates are plotted. (**C**) Average periods of clock oscillations in different neurons of *per*^L^ mutant flies. Indicated Gal4 drivers were used to activate LABL. Colors represent different temperatures as described in panel A. Bar represents the mean, +/- SD.

**Figure S2.**
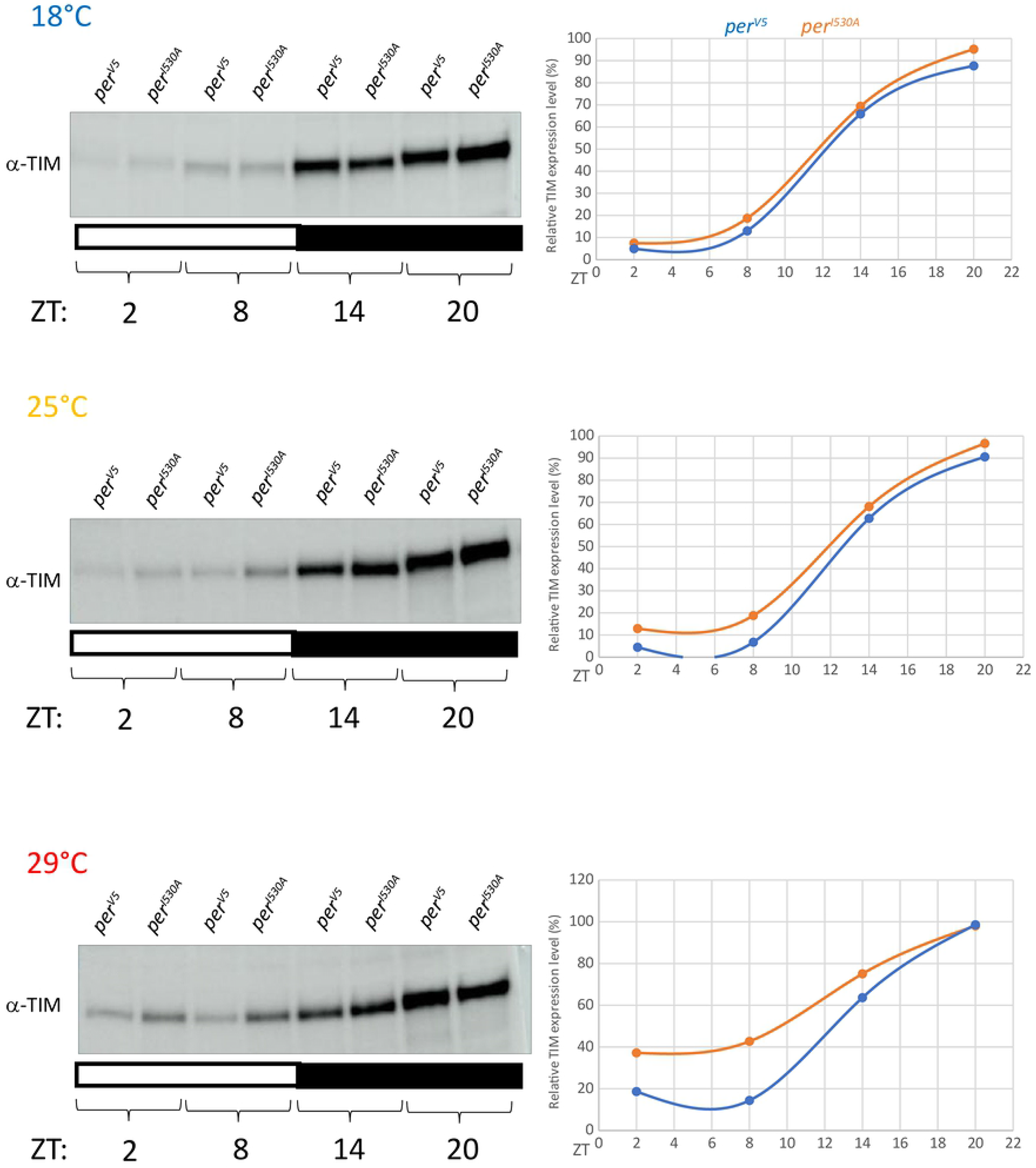
*per^I530A^* causes temperature-dependent dampening of TIM oscillations. Expression of TIM in *per^V5^* and *per^I530A-V5^* flies was compared at the indicated time points and temperatures. Left panel: Membranes that were initially blotted with anti-V5 antibodies (see Figure 3B) were stripped and developed with anti-TIM antibodies to analyze the effect of the PER mutation on TIM expression: Right panel: Relative TIM expression levels were plotted as line graphs for *per^V5^* and *per^I530-V5^* mutant flies. Error bars indicate SEM.

**Figure S3:**
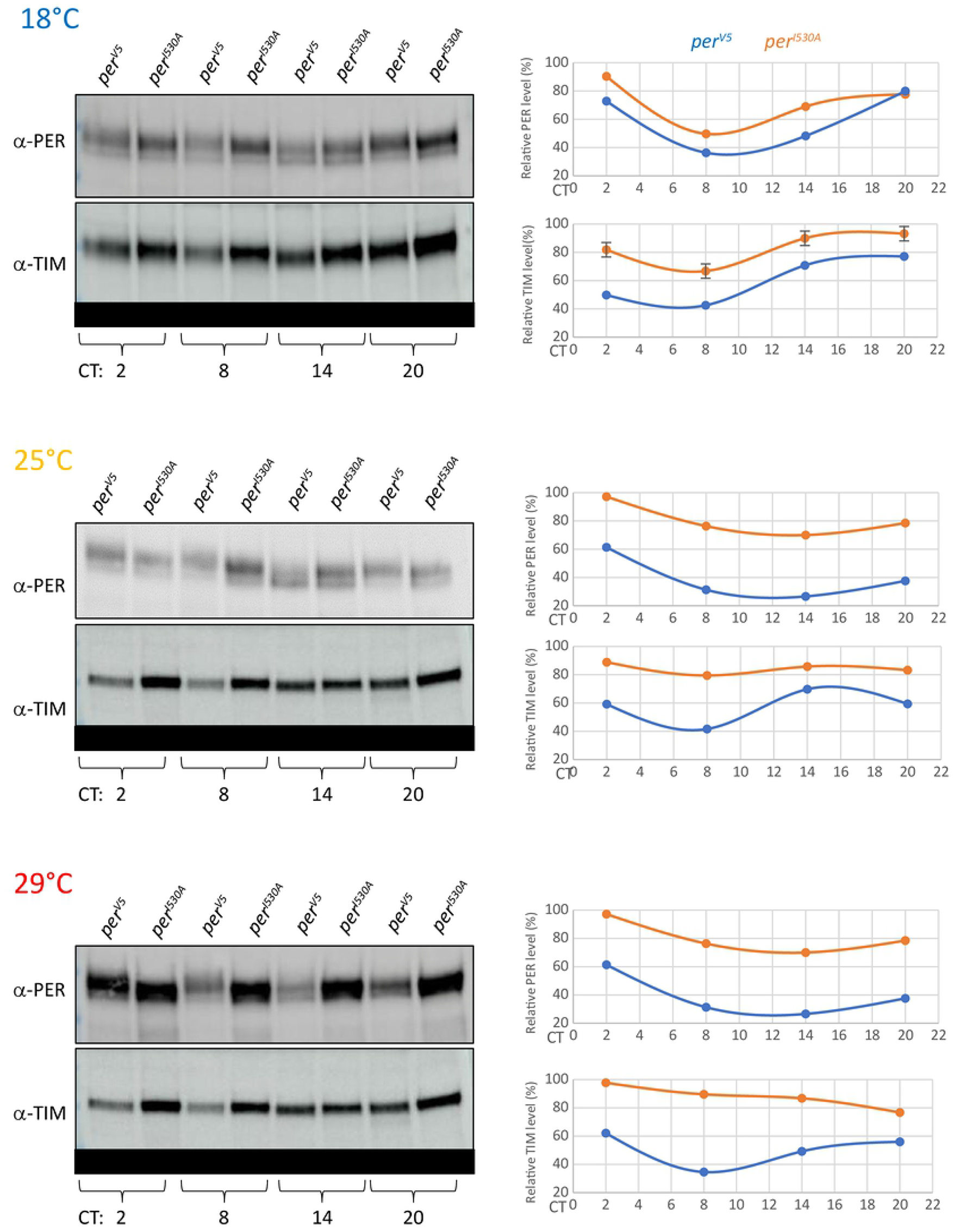
Analysis of PER and TIM expression in *per^V5^* and *per^I530A-V5^* fly heads at different temperatures in DD. Flies were kept for 3 days in LD and for 2 more days in DD at three constant temperatures (18°C, 25°C and 29°C). Left panels: Protein extracts from *per^V5^* and *per^I530A-V5^* heads were loaded next to each other for each time point to allow direct comparison. Blots were developed with anti-V5 or anti-TIM antibodies, respectively. Right panels: Relative expression levels were blotted as line graphs for *per^V5^* and *per^I530A-V5^* flies. Error bars indicate SEM.

**Figure S4:**
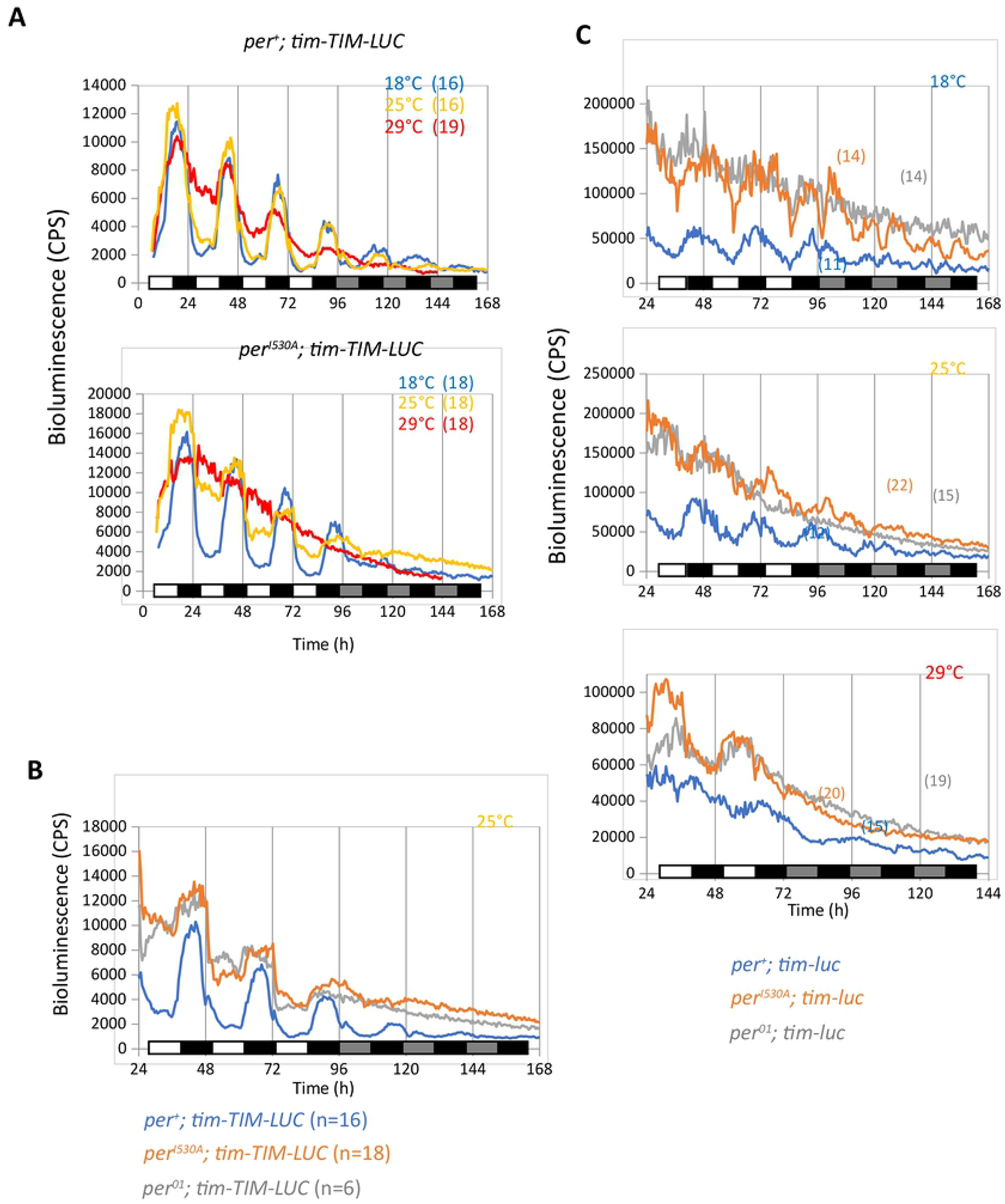
*per^I530A^* causes temperature-dependent dampening of TIM-LUC oscillations and impairs repressor function. **(A)** Bioluminescence rhythms of *per^+^* and *per^I530A^* flies expressing the *tim-TIM-LUC* fusion protein, reporting temporal TIM expression. Individual flies were measured at the indicated temperatures for 4 days in 12 hr : 12 hr LD, followed by 2 or 3 days in DD. White bars indicate lights on, black bars lights off, and grey bars subjective day. N numbers are indicated in parenthesis. **(B)** Bioluminescence rhythms of *per^+^*, *per^01^*, and *per^I530A^* flies expressing the *tim-TIM-LUC* fusion protein for 3 days in LD followed by 3 days in DD at 25°C. Labelling as in (A). **(C)** Bioluminescence rhythms of *per^+^*, *per^01^*, and *per^I530A^* flies expressing the *tim-luc* transgene, reporting *tim* transcription. Flies were recorded for 2 or 3 days in LD followed by 3 days in DD at the indicated temperatures. Labelling as in (A).

**Figure S5:**
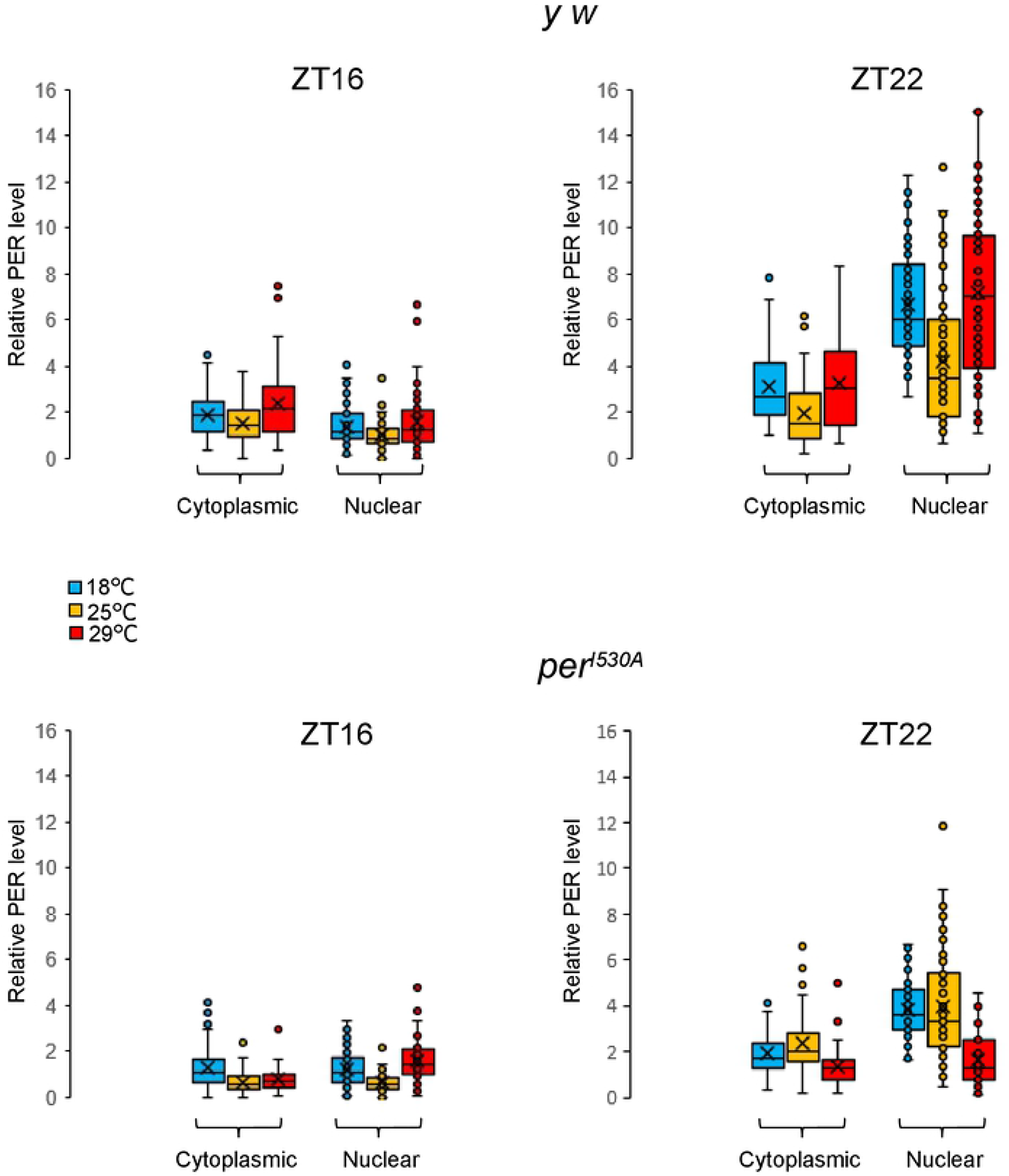
Relative PER level in the s-LNv in control flies and *per^I530A^* flies. Same cells as in Figure 6 were quantified. Note that in control flies PER levels appear reduced at ZT22 while the fraction of nuclear PER over cytoplasmic PER is constant at all three temperatures (Figure 6).

**Table S1:** Free-running behavioral rhythms of transgenic *per^01^; per^I530A^* and *per^01^; per^wild type^* flies

| Genotype | Temp (°C) | n | Tau (h ± SEM) | RS ± SEM | % rhythmic |
| --- | --- | --- | --- | --- | --- |
| *per<sup>01</sup>; 1-5-1/+; 2-2-2/+* (wild type) | 18 | 32 | 22.9 ± 0.3 | 1.9 ± 0.1 | 87.5 |
|  | 25 | 32 | 23.2 ± 0.1 | 1.9 ± 0.1 | 87.5 |
|  | 29 | 31 | 22.8 ± 0.1 | 2.6 ± 0.2 | 83.9 |
| *per<sup>01</sup>; 8-6; 7-3 (per<sup>l530A</sup>)* | 18 | 50 | 24.9 ± 0.4 | 1.9 ± 0.1 | 60.0 |
|  | 25 | 55 | 25.7 ± 0.3 | 2.5 ± 0.2 | 65.5 |
|  | 29 | 56 | 27.4 ± 0.3 | 1.6 ± 0.2 | 35.7 |

